# Periplasmic serine protease Prc is responsible for amyloid subunit CsgA degradation and proteostasis in *Escherichia coli*

**DOI:** 10.1101/2024.12.25.630354

**Authors:** Shinya Sugimoto, Yurika Terasawa, Naoki Tani, Kunitoshi Yamanaka, Yuki Kinjo

## Abstract

*Escherichia coli* synthesizes curli amyloid fibers extracellularly during biofilm formation and host colonization. The proteostasis network regulates the major curli subunit, CsgA, to prevent intracellular amyloid aggregation, yet the degradation mechanism remains elusive. Here, through a comprehensive investigation employing genetically engineered *E. coli*, multi-copy-suppressor screening, and biochemical analyses, we identify periplasmic serine protease Prc as a key player in CsgA degradation. Prc directly degrades CsgA through internal cleavage, differing from canonical tail-specific proteases. Although the bacterial HtrA homologs DegP and DegQ exhibit limited CsgA degradation activity in vitro in the presence of the suicide activator YjfN, deletion of these proteases did not affect native CsgA degradation in vivo. Instead, Prc, in coordination with the periplasmic chaperone CsgC, prevents the periplasmic accumulation of CsgA amyloid-like aggregates. Additionally, disruptions in secretion efficiency and proteolytic systems reduce *csg* operon expression through activation of the Rcs and Cpx two-component systems. These findings reveal a dual-layered strategy employed by *E. coli* to prevent intracellular accumulation of extracellular amyloids at both protein degradation and transcriptional regulation levels. This study provides insights into the mechanisms ensuring cellular homeostasis during curli biogenesis.

## Introduction

Proteins are the most versatile macromolecules in almost all cellular functions (*1*). They are synthesized on ribosomes as polypeptides and must fold into correct three-dimensional structures to perform their biological functions. However, protein folding is intrinsically error prone, and efficient protein folding throughout life is a tremendous challenge. The maintenance of protein homeostasis (proteostasis) to prevent accumulation of misfolded proteins and potentially toxic protein aggregates is a fundamental biological function. Loss of proteostasis associated with formation of toxic protein aggregates is linked to aging, various neurodegenerative diseases, and other pathologies affecting many different cell types (*2*). In particular, formation of amyloids, β-sheet-rich fibrillar aggregates of proteins and peptides, are deleterious for cells, as they are linked to severe human neurodegenerative diseases like Alzheimer’s (AD) and Parkinson’s (PD) diseases.

Despite their association with diseases, many organisms produce functional amyloids as a regular part of cellular biology, distinct from toxic amyloids as they serve specific biological functions (*3*). For example, curli, an extracellular amyloid fiber, is produced by enteric bacteria such as *Escherichia coli* and *Salmonella* species and plays a crucial role as a part of the extracellular matrix during biofilm formation. In *E. coli*, curli biogenesis requires seven curli-specific gene (*csg*) products (CsgA-G) (*4*). CsgA and CsgB are the major and minor structural components of curli, respectively. They are synthesized by ribosomes and transported from the cytoplasm to the periplasm (*4*). CsgA and CsgB are further secreted through the outer membrane by the nonameric bacterial amyloid secretion channel CsgG (*5–8*). The extracellularly secreted soluble CsgA binds CsgB, a nucleator for amyloid formation (*9*) and a scaffold binding to CsgF, forming a β-sheet structure leading to amyloid fibril formation on the cell surface (*10–12*).

An efficient secretion and chaperone network assures that CsgA does not form intracellular toxic amyloid aggregates. In the cytoplasm, molecular chaperone DnaK, in cooperation with its partners DnaJ and CbpA, prevents aggregation of CsgA and CsgB by recognizing their hydrophobic secretion signals, which promotes efficient translocation of CsgA to the periplasm (*13, 14*). In the periplasm, molecular chaperones CsgC (*15*), CsgE (*16*), and Spy (*17*) are thought to protect CsgA from aggregation, as they prevent amyloid formation of CsgA in vitro. However, the mechanisms governing intracellular degradation of CsgA remain largely unknown. Previous studies suggested that CsgA could be degraded by periplasmic proteases, yet the specific proteases responsible remained unidentified (*18*). Additionally, how *E. coli* cells respond to impaired proteostasis in the periplasm during curli biogenesis remains unclear.

In this study, we developed an analytical system to investigate the transient state of CsgA in the periplasm, elucidating its cleavage and aggregation dynamics. Our findings identify Prc as a key protease responsible for degrading CsgA in the periplasm. Prc collaborates with the periplasmic chaperone CsgC to ensure efficient degradation in the periplasm. Additionally, we discovered that in response to CsgA accumulation, negative feedback mechanisms downregulate the expression of *csg* genes.

## Results

### Evolutionarily conserved DegP/HtrA and its homolog DegQ are involved in CsgA- sfGFP degradation in the periplasm

The experimental detection of CsgA and CsgB within the periplasm has posed challenges due to their rapid transit during curli biogenesis, implying their transient presence in the periplasm of *E. coli* (*18*). To overcome this hurdle, we engineered a genetically modified *E. coli* strain for real-time monitoring of CsgA status in the periplasm. This strain, termed BA-FP, was designed to express CsgA-sfGFP and CsgB-mCherry from their native loci on the genome (Fig. 1A). The resulting BA-FP strain exhibited fluorescence signals of both CsgA-sfGFP and CsgB-Cherry at the cell periphery (Fig. 1B). As these fluorescent fusion proteins did not form foci, they are likely stabilized by molecular chaperones and/or degraded to the domains of fluorescent proteins by proteases.

**Fig. 1.**
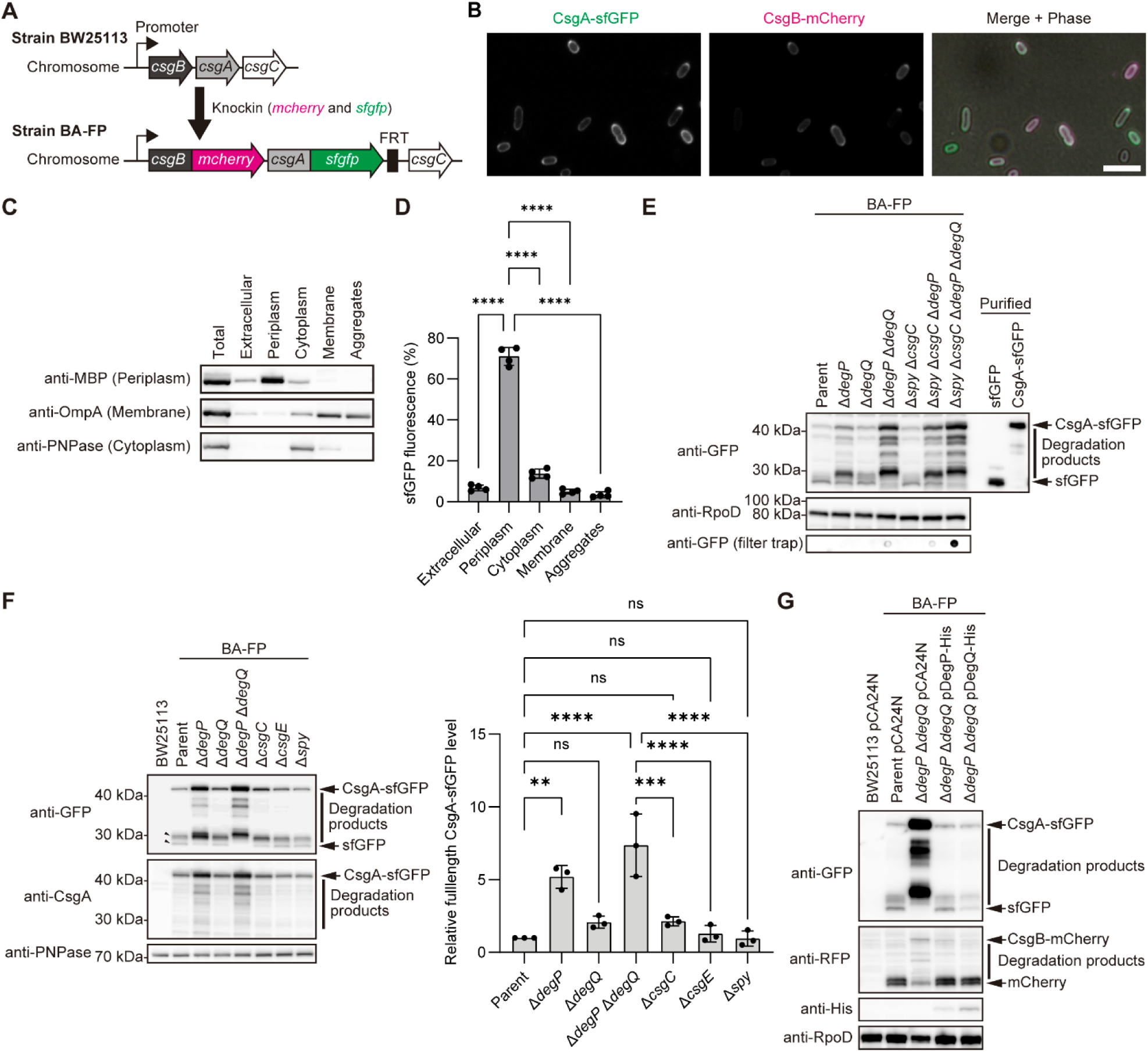
Involvement of DegP and DegQ in CsgA-sfGFP degradation in the periplasm. (**A**) Schematic illustration for construction of the reporter strain BA-FP expressing CsgA-sfGFP and CsgB-mCherry translational fusions from the chromosome of *E. coli* BW25113. FRT, a flippase recognition target site remained after removal of the Km^R^ cassette. (**B**) Fluorescence images of CsgA-sfGFP (green) and CsgB-mCherry (magenta) in the strain BA-FP. Scale, 5 μm. (**C**) Cell fractionation of the BA-FP strain was performed, and the resulting fractions were subjected to immunoblotting to detect MBP, OmpA, and PNPase, which serve as markers for periplasmic, cytoplasmic, and membrane proteins, respectively. (**D**) The relative fluorescence intensity of sfGFP was measured in the indicated fractions of BA-FP, with the total intensity of all fractions set to 100%. (**E**) Immunoblots of the indicated strains to detect CsgA-sfGFP. The bottom panel shows CsgA-sfGFP aggregation analyzed using filter-trap assay. (**F**) Effects of deletion of periplasmic protease genes on CsgA-sfGFP accumulation in the periplasm. CsgA-sfGFP was detected using immunoblotting. Band intensity of full-length CsgA-sfGFP were measured using LAS-4000 image analyzer. (**G**) Complementation of BA-FP Δ*degP* Δ*degQ* using the indicated plasmids. An empty plasmid (pCA24N) was used as a negative control. Parent pCA24N represents BA-FP transformed with pCA24N. Antibodies against GFP and CsgA were used to detect CsgA-sfGFP. Anti-RFP antibody was used to detect CsgB-mCherry. An anti-His-tag antibody was used to detect DegP-His and DegQ-His. RpoD and PNPase were detected as loading controls. Purified sfGFP and CsgA-sfGFP were used as positive controls. BW25113 was used as a negative control for immunoblotting for detection of fluorescent proteins. The positions of molecular mass markers are indicated at the left of the blots in E. Data are represented as mean ± SD from at least three independent experiments (one-way ANOVA followed by Tukey’s post hoc tests for D and F). \*\**P* < 0.01, \*\*\**P* < 0.001, \*\*\*\**P* < 0.0001. ns, not significant.

To demonstrate the localization of CsgA-sfGFP in the BA-FP strain, we performed cell fractionation. Immunoblotting for MBP, OmpA, and PNPase, markers for periplasmic, membrane, and cytoplasmic proteins respectively, confirmed successful separation of theses fractions, despite some observed contamination (Fig. 1C). Predominant fluorescence of sfGFP in the periplasmic fraction (Fig. 1D) indicated that CsgA-sfGFP was indeed expressed in the periplasm, but neither the cytoplasm nor the extracellular milieu.

Subsequently, immunoblotting with an anti-GFP antibody was conducted to detect CsgA- sfGFP. The analysis revealed significant cleavage of CsgA-sfGFP into sfGFP, with only a minor fraction of full-length CsgA-sfGFP detected (Fig. 1E, leftmost lane). Further immunoblotting using an anti-CsgA antibody unveiled predominant digestion of the N-terminal CsgA domain by proteases, as evidenced by decreased smaller bands reacting with the anti-CsgA antibody compared with those detected with anti-GFP antibody (Fig. 1F, arrowheads in the left second lane). Additionally, full-length CsgB-mCherry was scarcely detected in the periplasm of the BA-FP strain (Fig. 1G, left second lane). These findings suggest extensive cleavage of CsgA-sfGFP and CsgB-mCherry within the periplasm.

In humans, the widely conserved serine protease HtrA1 degrades various fragments of the amyloid β-precursor protein (*19*). In *E. coli*, three HtrA homologs, DegP, DegQ, and DegS, participate in protein quality control within the cell envelope (*20*). While DegP and DegQ are periplasmic, DegS serves as an inner-membrane-anchored protease vital for *E. coli* growth. Consequently, we investigated the potential involvement of non-essential DegP and DegQ in CsgA-sfGFP degradation within the periplasm. Deleting *degP* resulted in increased accumulation of full-length and partial degradation products of CsgA-sfGFP, while *degQ* deletion had a minor impact on CsgA-sfGFP degradation patterns (Fig. 1E). Simultaneous *degP* and *degQ* deletion further augmented the accumulation of full-length and degradation intermediates of CsgA-sfGFP (Fig. 1E). Complementation with plasmids carrying these genes restored CsgA-sfGFP degradation to parental strain levels (Fig. 1G), indicating a redundant role of DegP and DegQ in CsgA-sfGFP degradation, albeit with DegP exerting a more pronounced effect.

Despite several periplasmic chaperones, such as CsgC (*15*), CsgE (*16*), and Spy (*17*), known to prevent CsgA aggregation in vitro, their impact on CsgA degradation within the periplasm remained unclear. Deletion mutants of these genes generated using the BA-FP strain did not significantly affect CsgA-sfGFP degradation patterns (Fig. 1F). Only deletion of *csgC* marginally increased full-length CsgA-sfGFP accumulation (*p* = 0.505 by one-way ANOVA with Tukey’s test) (Fig. 1F). Analysis of combined deletion mutants showed no significant changes in the degradation patterns of CsgA-sfGFP. However, a filter-trap assay revealed increased amyloid-like SDS-insoluble aggregates of CsgA-sfGFP in mutants lacking Spy, CsgC, DegP, and DegQ compared to other mutants (Fig. 1E). These findings highlight the aggregation-prone nature of CsgA-sfGFP in vivo and suggest a potential interplay between periplasmic chaperones and proteases in stabilizing CsgA- sfGFP within the periplasm. Consistently, purified CsgA-sfGFP also exhibited an aggregation-prone behavior.

### Prc is also responsible for CsgA-sfGFP degradation in the periplasm

Although the full-length CsgA-sfGFP accumulated in the periplasm of the BA-FP Δ*degP* Δ*degQ* strain, it underwent cleavage at several sites (Fig. 1, E to G). Hence, we hypothesized that other proteases may be involved in CsgA-sfGFP degradation within the periplasm. To identify these proteases, we conducted multi-copy suppressor screening using the BW25113 Δ*degP* Δ*degQ* strain transformed with 21 selected plasmids from the ASKA library (*21*), which can express potential periplasmic proteases (https://www.uni-due.de/zmb/microbiology/e-coli-proteases.php#periplasm). In this experiment, BW25113 Δ*degP* Δ*degQ* was used as a host strain to identify DegP and DegQ-independent proteases that degrade native CsgA. Curli-negative colonies were selected on Congo Red (CR)- YESCA plates to amplify the inserted genes in the plasmid, and the amplified DNA was sequenced (Fig. 2, A to C, and fig. S1, A and B). This screening led to the identification of eight candidate periplasmic proteases (Fig. 2D).

**Fig. 2.**
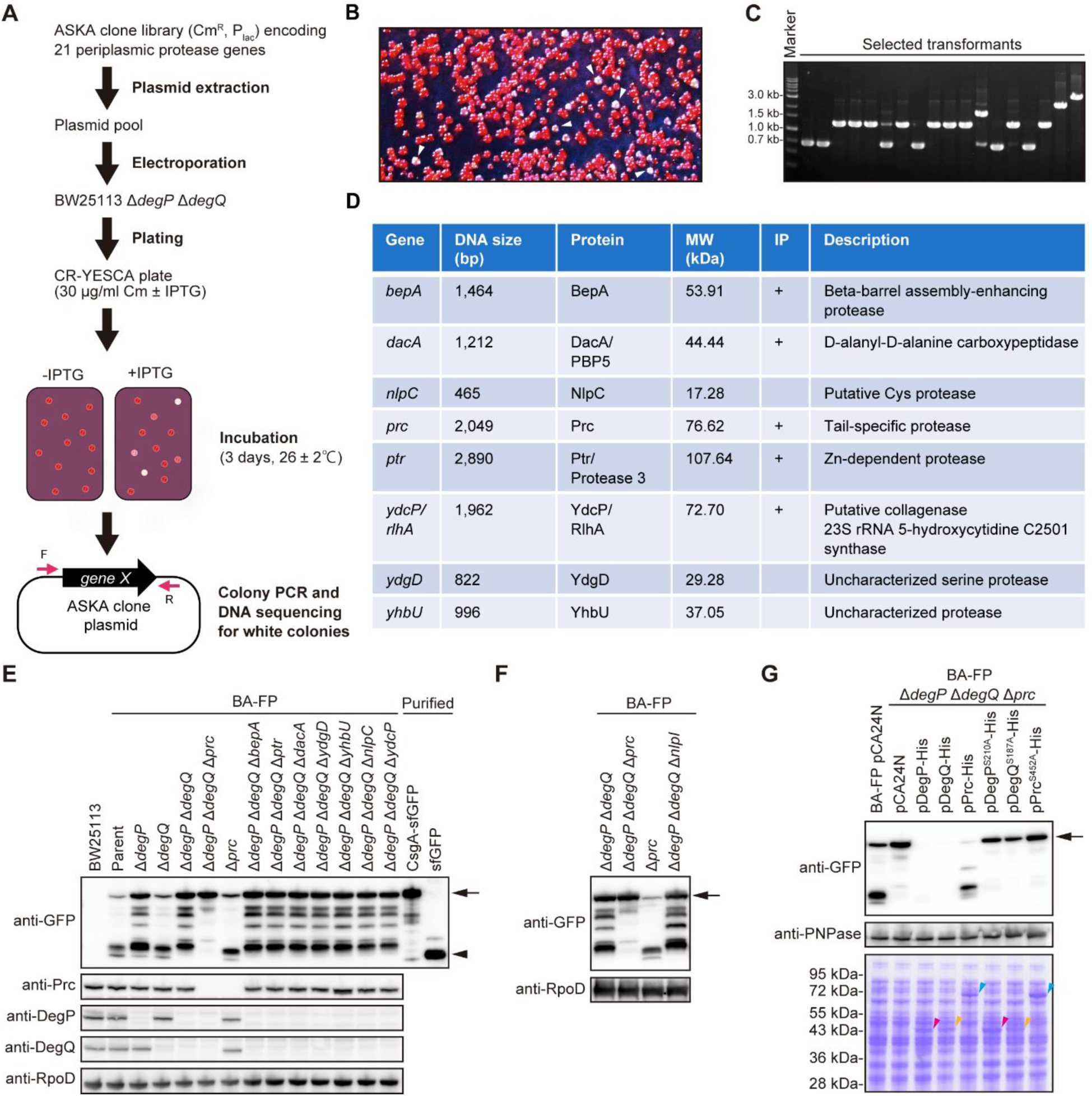
Identification of Prc as a protease responsible for CsgA-sfGFP degradation in the periplasm. (**A**) A schematic illustration for multi-copy suppressor screening to identify periplasmic proteases, other than DegP and DegQ, that are responsible for CsgA degradation. In this experiment, BW25113 Δ*degP* Δ*degQ* was used as a host strain to identify DegP and DegQ-independent proteases that degrade native CsgA. Magenta arrows indicate forward (F) and reverse (R) primers. (**B**) A typical picture of *E. coli* colonies grown on Congo Red-containing YESCA (CR-YESCA) plates supplemented with 0.01 mM IPTG. White arrowheads represent white colonies, candidate clones expressing a periplasmic protease involved in CsgA degradation. (**C**) Agarose gel electrophoresis for colony PCR products of the selected transformants. The amplified PCR products were purified and sequenced. (**D**) Candidates of periplasmic proteases involved in CsgA degradation in the periplasm identified using multi-copy suppressor screening. + indicates the CsgA- binding proteins identified using immunoprecipitation (IP) assay. (**E, F**) Immunoblotting to detect CsgA-sfGFP in the indicated strains. Prc, DegP, and DegQ were detected to confirm deletion in the respective genes. (**G**) Complementation of *degP*, *degQ*, and *prc* deletion in the strain BA-FP using plasmids expressing the wild-type and protease activity-defective mutant proteins. CsgA-sfGFP was detected using immunoblotting with anti-GFP antibody. RpoD and PNPase were detected as loading controls. The black arrows and arrowheads represent full-length CsgA-sfGFP and sfGFP, respectively. Magenta, yellow, and blue arrowheads mark DegP, DegQ, and Prc, respectively.

Additionally, we performed an immunoprecipitation assay in the presence of protease inhibitors to identify CsgA-binding proteins (fig. S2A). LC-MS/MS analysis identified 24 proteins, including known CsgA-binding proteins like CsgB, CsgC, CsgF, and CsgG (fig. S2B). Five candidate proteases (BepA, Ptr, DacA, YdcP, and Prc) were found in both experiments (Fig. 2D).

To determine the involvement of the identified potential periplasmic proteases in CsgA- sfGFP degradation, we deleted these genes from the BA-FP Δ*degP* Δ*degQ* strain. Among the generated strains, only the BA-FP Δ*degP* Δ*degQ* Δ*prc* strain showed nearly complete suppression of full-length CsgA-sfGFP degradation (Fig. 2E). The BA-FP Δ*degP* Δ*degQ* Δ*nlpI* strain displayed a degradation pattern for CsgA-sfGFP similar to that of the BA-FP Δ*degP* Δ*degQ* strain, suggesting that Prc did not rely on its adaptor protein, NlpI (*22*), for CsgA-sfGFP degradation (Fig. 2F, rightmost lane). Plasmid-borne DegP and DegQ degraded CsgA-sfGFP entirely in a protease activity-dependent manner, while Prc expression reduced full-length CsgA-sfGFP in a protease activity-dependent manner (Fig. 2G).

Furthermore, we generated single-gene deletions in the BA-FP strain. The BA-FP Δ*prc* strain exhibited a CsgA-sfGFP band pattern similar to the BA-FP Δ*degQ* strain (Fig. 2E). In contrast, single deletions of the other candidate protease genes (*bepA*, *ptr*, *dacA*, *ydgD*, *yhbU*, *nlpC*, and *ydcP*) in the BA-FP strain did not affect the CsgA-sfGFP degradation pattern (fig. S3).

In summary, our multi-copy suppressor screening, combined with gene deletion analysis, identified the tail-specific protease Prc as a key periplasmic protease responsible for CsgA-sfGFP degradation within the periplasm.

### DegP and DegQ degrade unfolded, unmodified CsgA in a suicide activator-dependent mechanism

We explored whether the identified periplasmic proteases could degrade unfolded CsgA lacking the sfGFP domain. DegP and DegQ did not exhibit degradation activity towards non-tagged CsgA (Fig. 3, A and B), C-terminal His-tagged CsgA (CsgA-His, fig. S4, A and B), or N-terminal His-tagged CsgA (His-CsgA, fig. S4C). As DegP sometimes relies on adaptor or suicide activator proteins to degrade specific substrates, we investigated the influence of the the adaptor CpxP (*23*) and suicide activator YjfN (*24*) on His-CsgA degradation by DegP. Notably, DegP degraded His-CsgA in the presence of YjfN but not CpxP (fig. S4, C to E). Similarly, DegQ required YjfN for the degradation of His-CsgA (fig. S4E). Furthermore, unmodified CsgA was susceptible to degradation by DegP and DegQ, only in the presence of YjfN, with DegP-YjfN exhibiting higher activity towards CsgA compared to DegQ-YjfN (Fig. 3, A and B).

**Fig. 3.**
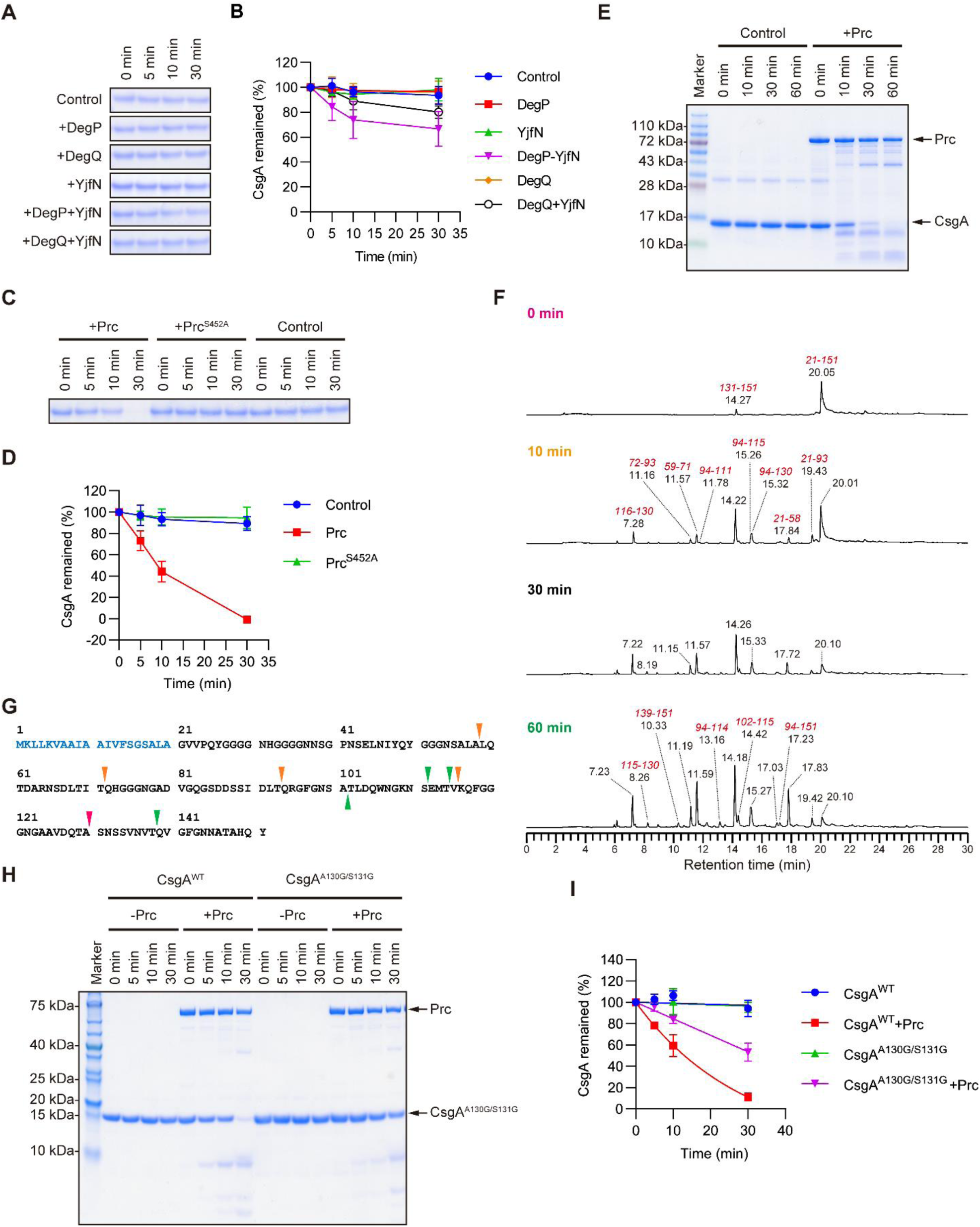
In vitro degradation of unfolded CsgA by the periplasmic proteases. (**A**) Degradation of purified, unfolded CsgA (5 μM) by purified DegP or DegQ (5 μM) in the absence and presence of YjfN (25 μM) was analyzed using SDS-PAGE. (**B**) The band intensity of CsgA in (A) was quantified using the ImageQuant TL software. The intensity at 0 min was set as 100%. Data are represented as mean ± SD from at least four independent experiments. (**C**) Degradation of purified CsgA (5 μM) by purified Prc or its protease activity-deficient mutant (Prc^S452A^) (5 μM) was analyzed using SDS-PAGE. (**D**) The band intensity of CsgA in (C) was quantified as in (B). (**E**) For LC-MS/MS analysis, to identify digestion sites of Prc in CsgA, purified, unfolded CsgA (50 μM) was degraded by Prc (5 μM), and the degradation products were analyzed using SDS-PAGE. (**F**) LC-MS/MS analysis for time-dependent fragmentation of CsgA by Prc. The assigned peaks at the indicated retention times on the chromatogram of LC correspond to CsgA peptides with unique amino acid numbers colored in red. (**G**) Prc-digestion sites in CsgA identified using LC-MS/MS. Blue characters represent the secretion signal. Arrowheads indicate Prc-digestion sites detected at 0 (magenta), 10 (yellow), and 60 min (green). (**H**) Degradation of purified, unfolded CsgA^A130G/S131G^ (5 μM) by purified Prc (5 μM) was analyzed using SDS-PAGE. (**I**) The band intensity of CsgA in (H) was quantified as in (B).

### Prc directly degrades unfolded, unmodified CsgA by a mechanism distinct from that of canonical tail-specific proteases

We investigated whether Prc could degrade unfolded, unmodified CsgA in vitro. Unlike DegP and DegQ, Prc exhibited the ability to degrade unfolded, unmodified CsgA on its own, while a protease activity-deficient mutant, Prc^S452A^, did not (Fig. 3, C and D). Additionally, Prc effectively degraded unfolded CsgA-His (fig. S4, A and B). Notably, the degradation rate of unfolded, unmodified CsgA by Prc was significantly faster compared to DegP-YjfN and DegQ-YjfN (Fig. 3, A to D).

Traditionally identified as a tail-specific protease, Prc is known to recognize and cleave the C-terminal tail region of substrate proteins, such as penicillin-binding protein 3, to facilitate protein maturation (*25*), as well as the SsrA-tag of improperly translated proteins (*26*) and proteins with nonpolar carboxy-terminal tags (*27*). Interestingly, we discovered that Prc could degrade unfolded, unmodified CsgA lacking a nonpolar carboxy-terminal tag (Fig. 3, C and D), as well as CsgA-sfGFP (Fig. 2G), and CsgA-His (fig. S4, A and B). On the other hand, Prc did not degrade aggregated CsgA and curli fibrils (Sugimoto *et al*. personal observation), indicating that Prc act on CsgA prior to amyloid formation.

Sodium dodecyl sulphate-polyacrylamide gel electrophoresis (SDS-PAGE) analysis revealed that Prc cleaved unfolded CsgA at several specific sites (Fig. 3E), indicating a mode of action distinct from canonical tail-specific proteases. To explore deeper into the cleavage sites of Prc in CsgA, we employed liquid chromatography-mass spectrometry (LC-MS). The LC chromatogram illustrated a time-dependent cleavage pattern of unfolded CsgA by Prc, and the peaks were identified using MS/MS analysis (Fig. 3F and fig. S5). A total of 9 Prc cleavage sites in CsgA were mapped to the internal region (Fig. 3G). Notably, the peak corresponding to the fragment of 131-151 appeared immediately after the addition of Prc (t = 0 min), suggesting the initial cleavage occurred at the peptide bond between Ala-130 and Ser-131. To understand the significance of this primary cleavage in subsequent digestion of CsgA, we utilized a CsgA mutant where Ala-130 and Ser-131 were substituted with Gly (CsgA^A130G/S131G^). Prc degraded unfolded CsgA^A130G/S131G^ similarly to wild type unfolded CsgA (CsgA^WT^) (Fig. 3H), albeit at a slightly slower rate than unfolded CsgA^WT^ (Fig. 3I). This implies that while the initial cleavage at the peptide bond between Ala-130 and Ser-131 enhances the efficiency of unfolded CsgA degradation, it is not indispensable for the subsequent cleavage at other sites.

In summary, our findings reveal a non-canonical mechanism of action of Prc, wherein it cleaves a substrate protein in the internal region rather than the C-terminal tail.

### Prc degrades unmodified CsgA in cells

We investigated the dynamics of unmodified/native CsgA in a wild-type background. Single and combined deletions of *degP*, *degQ*, and *prc* did not affect the total amount of CsgA that polymerized on the surface of the cell (Fig. 4A, +hexafluoroisopropanol, HFIP). However, in the Δ*prc* strain, CsgA was detectable without HFIP treatment (Fig. 4A, - HFIP). Previous reports suggested that Prc is a periplasmic protein associated with the cytoplasmic membrane (*25*); however, Singh et al. found that Prc is a soluble periplasmic protein (*22*). These results suggest the transient presence of non-amyloid CsgA in the periplasm of the Δ*prc* strain.

**Fig. 4.**
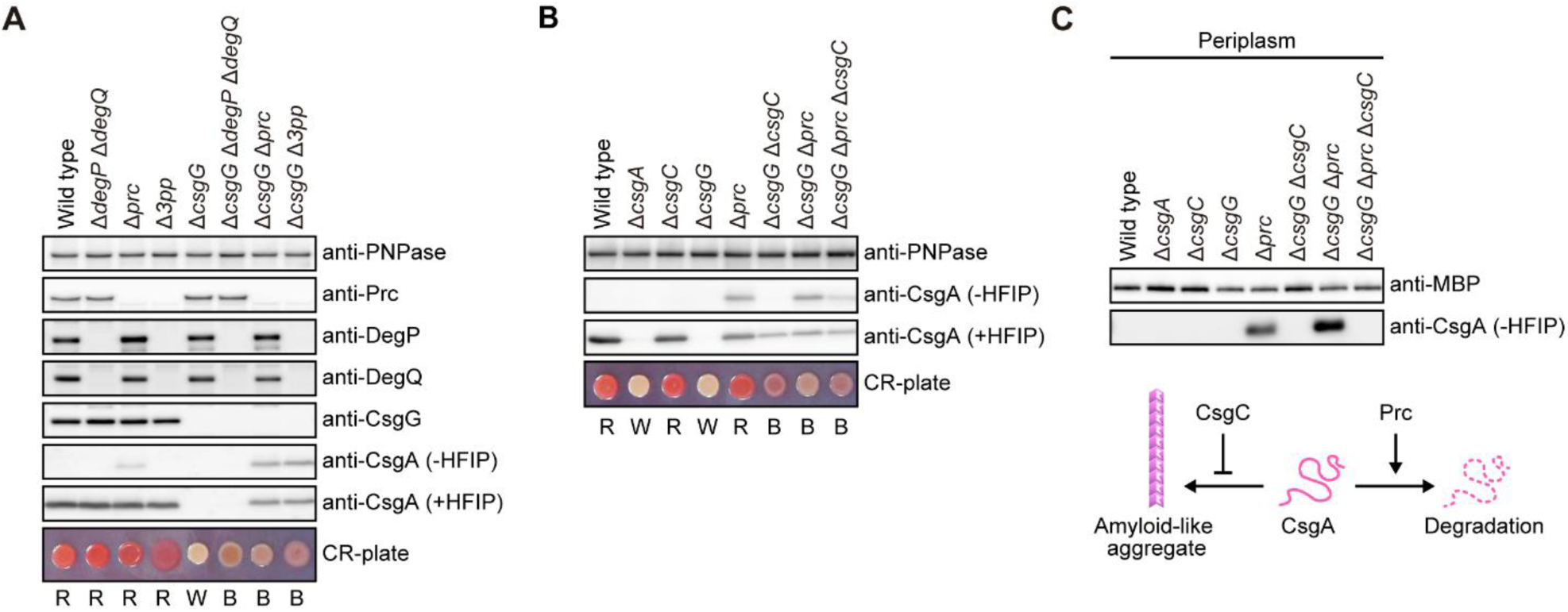
Cooperation of Prc and CsgC in quality control of CsgA in the periplasm. Immunoblotting to detect CsgA in the indicated strains using whole cells (**A, B**) and the periplasmic fractions (**C**). Δ*3pp*, deletion of 3 periplasmic protease genes (*degP*, *degQ*, and *prc*). HFIP was used to depolymerize curli to CsgA monomers (+HFIP). PNPase was detected as a loading control, while MBP was used as a marker for periplasmic proteins. Colony colors R, W, and B represent red, white, and brown, respectively.

To further explore CsgA accumulation in the periplasm, we deleted *csgG* from the Δ*prc*, Δ*degP* Δ*degQ*, and Δ*degP* Δ*degQ* Δ*prc* strains (referred to as Δ*3pp*, lacking three periplasmic protease genes). Consistent with findings by Loferer et al. (*18*), CsgA was undetectable in the Δ*csgG* strain via immunoblotting (Fig. 4A). Despite the Δ*csgG* Δ*degP* Δ*degQ* strain forming brown colonies on Congo Red plates, CsgA was not detected by immunoblotting, likely due to expression levels being below the detection threshold (Fig. 4A). However, SDS-soluble CsgA was detected in the Δ*csgG* Δ*prc* and Δ*csgG* Δ*3pp* strains (Fig. 4A, -HFIP), similar to the Δ*prc* strain.

These results suggest that Prc degrades unmodified CsgA in the periplasm of both wild-type and Δ*csgG* strains.

### Prc cooperates with CsgC to prevent accumulation of CsgA amyloid-like aggregates in the periplasm

The periplasmic chaperone CsgC is known to prevent amyloid formation of CsgA both in vitro and in vivo (*15*). To investigate the cooperation between CsgC and Prc in CsgA quality control, we performed a series of mutation experiments. Consistent with the previous report (*15*), SDS-insoluble CsgA was detected in the Δ*csgG* Δ*csgC* strain via immunoblotting (Fig. 4B, +HFIP), suggesting the accumulation of CsgA amyloid-like aggregates in the periplasm of this mutant. Notably, the level of SDS-soluble CsgA was reduced in the Δ*csgG* Δ*prc* Δ*csgC* strain compared to the Δ*csgG* Δ*prc* strain (Fig. 4B, - HFIP), despite the total amounts of CsgA being similar in both strains (Fig. 4B, +HFIP). These results suggest that Prc cooperates with CsgC to prevent accumulation of CsgA amyloid-like aggregates in the periplasm.

To further examine CsgA accumulation in the periplasm, we performed immunoblotting on isolated periplasmic fractions from the indicated strains, following the procedure used in Fig. 1C. CsgA was detected in the periplasmic fraction of the Δ*prc* and Δ*csgG* Δ*prc* strains but not the Δ*csgG* Δ*prc* Δ*csgC* strain (Fig. 4C). This indicates that CsgA exits in a soluble state in the periplasm of the Δ*prc* and Δ*csgG* Δ*prc* strains, and that CsgC prevents CsgA aggregation in the periplasm. Additionally, more CsgA was observed in the periplasm of the Δ*csgG* Δ*prc* strain compared with the Δ*prc* strain, which is expected as the deletion of *csgG* blocks the transport of CsgA from the periplasm to the extracellular environment.

These findings indicate that Prc is one of the proteases involved in degradation of native CsgA in the periplasm and that CsgC maintains CsgA in a state susceptible to targeting by Prc (Fig. 4C, lower).

### The Rcs and Cpx pathways negatively regulate csg gene expression in response to CsgA accumulation in the periplasm

We anticipated that in the absence of CsgG and the proteases involved in CsgA degradation, CsgA would accumulate in the periplasm. However, immunoblot analysis revealed that CsgA protein levels in the Δ*csgG* Δ*prc* and Δ*csgG* Δ*3pp* strains were lower than those in the wild-type and Δ*prc* strains (Fig. 4A). This suggests that CsgA expression might be reduced at the transcriptional level in these strains. To validate this hypothesis, we measured the mRNA levels of *csgA* using quantitative real-time polymerase chain reaction (qRT-PCR). The results showed significantly lower levels of *csgA* mRNA in the Δ*3pp*, Δ*csgG* Δ*prc*, and Δc*sgG* Δ*3pp* strains compared to the wild-type strain (*p* < 0.001) (Fig. 5A). Moreover, the *csgA* mRNA levels in the Δ*csgG* Δ*prc* and Δ*csgG* Δ*3pp* strains were lower than those in the Δ*prc* and Δ*csgG* strains (Fig. 5A). These findings indicate that disturbances in periplasmic protein degradation and secretion simultaneously reduce CsgA expression at the transcriptional level.

**Fig. 5.**
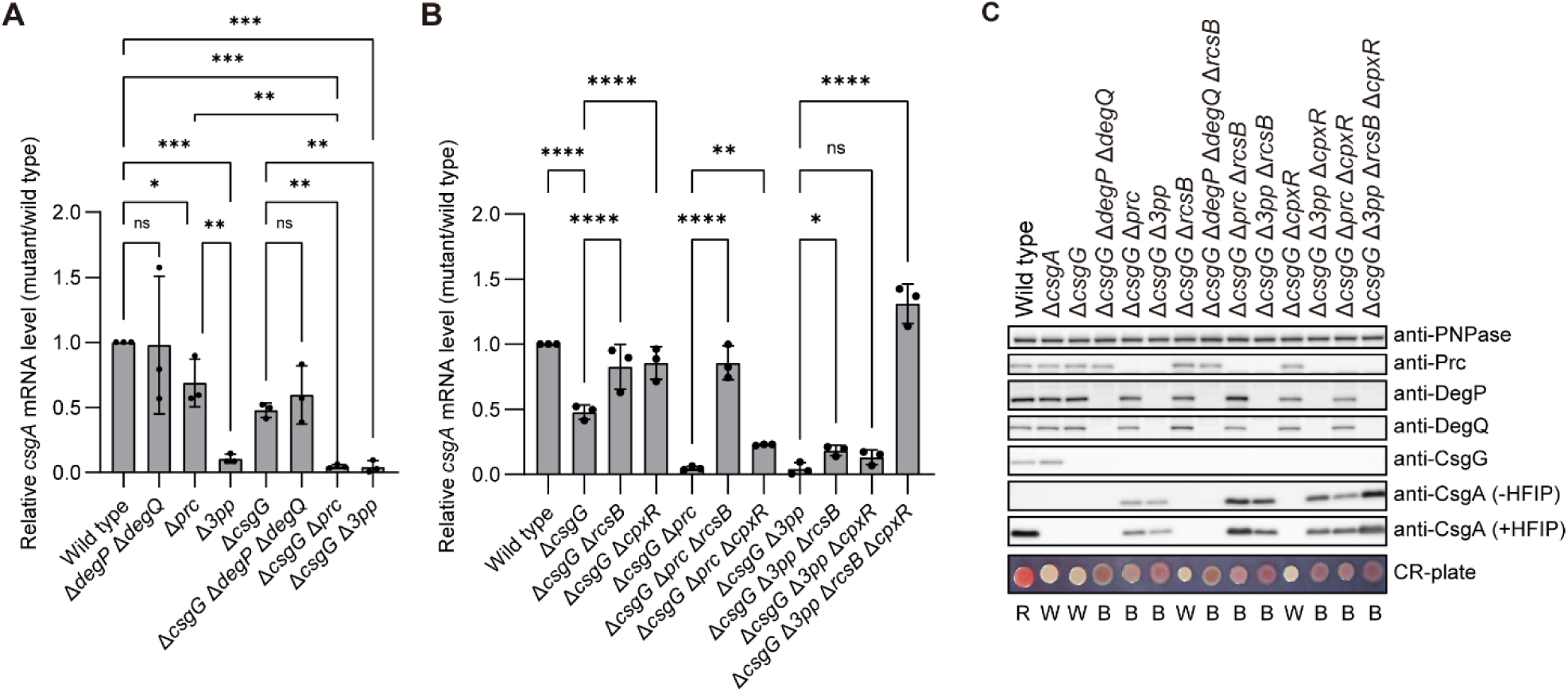
Downregulation of transcription of *csgA* via the response regulators of the Rcs and Cpx two component systems. (**A, B**) The relative mRNA levels of *csgA* in the indicated strains were analyzed using qRT-PCR. The mRNA levels in the wild-type strain were defined as 1. Δ*3pp*, deletion of 3 periplasmic protease genes (*degP*, *degQ*, and *prc*). (**C**) Immunoblotting to detect CsgA in the indicated strains. HFIP was used to depolymerize curli to CsgA monomers (+HFIP). PNPase was detected as a loading control. Colony colors R, W, and B represent red, white, and brown, respectively. Data are represented as mean ± SD from three independent experiments (one-way ANOVA with two-stage linear step-up procedure of Benjamini, Krieger, and Yekutieli correction with a False Discovery Rate of 5% for A and B). \**P* < 0.05, \*\**P* < 0.01, \*\*\**P* < 0.001, \*\*\*\**P* < 0.0001. ns, not significant.

Both the Rcs and Cpx two-component pathways are known to negatively regulate *csgBAC* and *csgDEFG* operon expression in response to changes in the cell envelope and membrane integrity (*28, 29*). Additionally, the Cpx pathway is activated in response to the accumulation of misfolded and aggregated periplasmic proteins, leading to the upregulation of various periplasmic chaperones and proteases (*30*). Therefore, we investigated the involvement of two transcriptional regulators, RcsB and CpxR, in the negative feedback regulation of the *csg* operons in response to CsgA accumulation in the periplasm. Deletion of *rcsB* reversed the reduced expression level of *csgA* mRNA observed in the Δ*csgG* Δ*prc* strain to that of the wild-type strain, while simultaneous deletion of *rcsB* and *cpxR* improved the expression level of csgA mRNA in the Δ*csgG* Δ*3pp* strain (Fig. 5B).

Finally, we confirmed the CsgA protein levels in mutants lacking RcsB and/or CpxR using immunoblotting. The CsgA protein levels in the Δ*csgG* Δ*prc* and Δ*csgG* Δ*3pp* strains were elevated upon deletion of *rcsB* and *cpxR* (Fig. 5C). Overall, the immunoblotting data were highly consistent with the qRT-PCR results.

These results indicate that the Rcs pathway primarily mediates the negative feedback regulation of the *csgBAC* operon in response to CsgA accumulation in the periplasm, with the Cpx pathway serving as a backup system that readily responds to such conditions.

## Discussion

The mechanisms governing the quality control of bacterial functional amyloids within the periplasm have long remained elusive. In this study, we revealed that Prc, previously recognized as a tail-specific protease, serves as a key periplasmic protease responsible for degrading CsgA independently of adaptors. To our knowledge, this marks the first identification of periplasmic proteases accountable for degrading the major subunit of bacterial functional amyloids in this compartment.

Building upon our discoveries and preceding groundbreaking investigations (*4, 7, 15, 18*), we propose several mechanisms for maintaining proteostasis in response to CsgA accumulation within the periplasm. Under normal conditions (the wild-type strain), CsgA is translocated from the cytoplasm to the periplasm via the Sec translocon and subsequently secreted through the CsgG channel into the extracellular environment. During curli biosynthesis, only a fraction of CsgA undergoes degradation by Prc in the periplasm, potentially regulating the number of curli fibers. Conversely, when efficient CsgA secretion is hindered by *csgG* deletion, Prc primarily targets CsgA degradation in the periplasm. Notably, our findings indicate that the periplasmic chaperone CsgC collaborates with the protease Prc to efficiently degrade CsgA in this compartment. In the strains lacking *csgG*, *prc*, *degP*, and *degQ* simultaneously (the Δ*csgG* Δ*3pp* strain), the accumulation of CsgA and other misfolded proteins in the periplasm may induce heightened periplasmic stress, thereby triggering a more robust negative feedback regulation of *csg* gene expression via the Rcs and Cpx pathways. Our results highlight the predominant role of the Rcs pathway in the negative feedback regulation of the *csg* operon in response to CsgA accumulation, with the Cpx pathway serving as a supplementary mechanism.

To screen for periplasmic proteases that degrade CsgA in vivo, we engineered and utilized the *E. coli* strain, BA-FP, which expresses CsgA-sfGFP from its genome. This artificial system proved effective in monitoring the localization and degradation patterns of CsgA- sfGFP. Through this approach, we identified three periplasmic serine proteases—DegP, DegQ, and Prc—that degrade CsgA-sfGFP in the periplasm. However, our results highlight the differing contributions of these proteases across experimental setups. Specifically, *degP* deletion had a greater impact on CsgA-sfGFP degradation in the BA- FP strain, while only *prc* deletion affected native CsgA degradation in the wild-type background. In vitro experiments further demonstrated that Prc, rather than DegP and DegQ, effectively degraded unmodified CsgA (Fig. 3, A to D). These findings suggest that investigations using a wild-type background are more relevant for understanding protein roles under physiological conditions, serving as a cautionary note for similar studies. Additionally, we can not rule out the involvement of other proteases in CsgA degradation in the periplasm. Notably, CsgA-sfGFP degradation persisted slightly in the BA-FP Δ*degP* Δ*degQ* Δ*prc* strain (Fig. 2E), indicating that other proteases, such as DegS, might also contribute.

To identify proteases responsible for CsgA degradation in the periplasm, we conducted multi-copy suppressor screening using the BW25113 Δ*degP* Δ*degQ* strain transformed with 21 selected plasmids from the ASKA library (*21*). Through selection of Curli-negative colonies on CR-YESCA plates, we successfully identified Prc as a key protease. For efficiency, we electroporated a plasmid pool containing all 21 plasmids instead of individual plasmids. This approach significantly saved time and costs associated with plasmid purification and electroporation. When conducting similar experiments with a larger number of plasmids, using a plasmid pool would be more advantageous compared to handling individual plasmids.

Protein quality control systems are indispensable for diverse cellular activities and are expected to be evolutionarily conserved across life forms. In a recent study, we demonstrated that the DnaK chaperone system, a bacterial Hsp70 system, prevents CsgA aggregation in the cytoplasm of *E. coli* (*13*), similar to the protective role of human Hsp70 in preventing Aβ_42_ aggregation (*31*). Moreover, mutations in DNAJC6/Hsp40, a co-chaperone of Hsp70, were linked to early-onset Parkinson’s disease in humans (*32*). Therefore, the Hsp70 chaperone systems play an important protective role for life by preventing toxic amyloid aggregation not only in bacteria but also in humans. Previous report indicated that human HtrA1 directly degrades Aβ_42_ (*19*). These findings underscore the critical protective function of Hsp70 chaperone systems against toxic amyloid aggregation, spanning from bacteria to humans. Additionally, while human HtrA1 was previously shown to directly degrade Aβ_42_, our study reveals that the evolutionarily conserved DegP and DegQ degrade CsgA in a YjfN-dependent manner in vitro. These observations suggest that proteases involved in degrading amyloid precursor proteins are evolutionarily conserved from bacteria to humans. However, the requirement of activator proteins like YjfN for efficient degradation by human HtrA1 remains to be elucidated, except for Tau, which is associated with neurodegenerative diseases and is degraded by HtrA1 alone (*33*). Furthermore, investigating the molecular evolution of Prc, the primary protease responsible for CsgA degradation in the periplasm, may shed light on amyloid-regulating proteases’ evolution and offer valuable insights into understanding and treating human neurodegenerative diseases linked to toxic amyloid aggregates.

## Materials and Methods

### Strain construction

The *E. coli* BA-FP (Table S1) was constructed as follows. First, the *csgA-sfgfp* translational fusion gene was inserted into the BW25113 genome. BW25113 was transformed with pKD46 and transformants were selected on Lysogeny Broth (LB) agar plates containing 100 μg/mL Ampicillin (Amp) at 30°C. The transformants were cultured at 30°C in SOB medium (2% tryptone, 0.5% yeast extract, 10 mM NaCl, 2.5 mM KCl, 20 mM MgSO_4_) containing 100 μg/mL Amp and 0.2% arabinose. When optical density of the culture reached 0.5, bacterial cells were harvested by centrifugation at 5,000 ×g for 10 min at 4°C and then washed three times with distilled and deionized water. The cells were suspended in 10% glycerol solution and used for electroporation. The chimeric DNA fragment containing partial 3’-end of *csgB* fused to the mCherry gene and entire *csgA*- fused to the sfGFP gene was amplified by PCR using KOD Plus Neo DNA polymerase (Toyobo, Osaka, Japan), primers CsgB-mCherry-SOE-F and sfGFP-Km-SOE-R, and pBAD-CsgB-mCherry-CsgA-sfGFP (Table S2) as a template. The DNA fragment containing FRT-Km^R^-FRT connected to the 20-bp downstream region of *csgA* was amplified by PCR using KOD Plus Neo DNA polymerase, primers sfGFP-Km-SOE-F and Km-CsgA-3’-SOE-R (Table S2), and pKD4 as a template. These DNA fragments were connected and amplified by splicing overlapping elongation-polymerase chain reaction (SOEing-PCR) using KOD Plus Neo DNA polymerase and the primers CsgB-SOE-F3 and CsgA-3’-R (Table S2). The resulting SOEing-PCR fragment was purified using a QIAquick PCR purification kit (Qiagen, Tokyo, Japan) and transformed into competent BW25113 cells harboring pKD46 by electroporation using a MicroPulser (BioRad, Hercules, CA, USA). Kanamycin (Km)-resistant (Km^R^) colonies were selected on LB agar plates containing 50 μg/mL Km at 37°C, and gene integration into the chromosome was checked using colony PCR, followed by DNA sequencing. The Km^R^ cassette sandwiched by FRT was removed from the chromosome using pCP20 (*34*). Competent cells of BA-FP cells harboring pKD46 were prepared as described above and used for further deletion of genes of interest.

BA-FP deletion mutants (Δ*degP*, Δ*degQ*, Δ*degP* Δ*degQ*, Δ*spy* Δ*csgC*, Δ*spy* Δ*csgC* Δ*degP* Δ*degQ*, Δ*prc*, Δ*degP* Δ*degQ* Δ*prc*, Δ*degP* Δ*degQ* Δ*bepA*, Δ*degP* Δ*degQ* Δ*ptr*, Δ*degP* Δ*degQ* Δ*dacA*, Δ*degP* Δ*degQ* Δ*ydgD*, Δ*degP* Δ*degQ* Δ*yhbU*, Δ*degP* Δ*degQ* Δ*nlpC*, Δ*degP* Δ*degQ* Δ*ydcP*, Δ*degP* Δ*degQ* Δ*nlpI*, Δ*bepA*, Δ*ptr*, Δ*dacA*, Δ*ydgD*, Δ*yhbU*, Δ*nlpC*, and Δ*ydcP*) were constructed using the one-step method for inactivation of chromosomal genes (*34*). The plasmids and primers used for gene knockout are listed in Tables S1 and S2, respectively. The RFT-Km^R^-RFT genes sandwiched between the upstream 100 bp and downstream 100 bp fragments of the respective genes, except for *prc*, were amplified from the respective mutant strains of the KEIO collection (*35*) using PCR with the respective primer sets (Table S2). As we found that JW1819-KC distributed to us was not successfully constructed, the RFT-Km^R^-RFT gene sandwiched between the 100 bp upstream and downstream regions of *prc* was amplified using PCR with KOD Plus polymerase, primers prc-KO-F1 and prc-KO-R1, and pKD4 as a template. The amplified DNA fragments were transformed into competent BA-FP cells harboring pKD46, and KmR transformants were screened on LB agar plates containing 50 μg/mL Km at 37°C. Gene deletion was checked using colony PCR and DNA sequencing.

The KEIO collection was used as a single-gene deletion mutant of BW25113, except for *prc*. Therefore, we constructed BW25113 Δ*prc* as mentioned above. Multiple-gene deletion mutants of BW25113 (Δ*degP* Δ*degQ*, Δ*degP* Δ*degQ* Δ*prc*, Δ*csgG* Δ*degP* Δ*degQ*, Δ*csgG* Δ*prc*, Δ*csgG* Δ*degP* Δ*degQ* Δ*prc*, Δ*degP* Δ*degQ* Δ*prc* Δ*rcsB*, Δ*degP* Δ*degQ* Δ*prc* Δ*cpxR*, Δ*degP* Δ*degQ* Δ*prc* Δ*rcsB* Δ*cpxR*, Δ*rcsB* Δ*cpxR*, Δ*csgG* Δ*csgC*, Δ*csgG* Δ*prc*, Δ*csgG* Δ*prc* Δ*csgC*, Δ*csgG* Δ*rcsB*, Δ*csgG* Δ*degP* Δ*degQ* Δ*rcsB*, Δ*csgG* Δ*prc* Δ*rcsB*, Δ*csgG* Δ*degP* Δ*degQ* Δ*prc* Δ*rcsB*, Δ*csgG* Δ*cpxR*, Δ*csgG* Δ*degP* Δ*degQ* Δ*prc* Δ*cpxR*, Δ*csgG* Δ*prc* Δ*cpxR*, and Δ*csgG* Δ*degP* Δ*degQ* Δ*prc* Δ*rcsB* Δ*cpxR*) were also constructed following the same procedure (Table S1).

### Plasmid construction

The DNA fragment containing *csgA* with its stop codon was amplified from the genomic DNA of BW25113 using PCR with KOD Plus DNA polymerase and the primer set His-CsgA-pET28-Nde-F and His-CsgA-pET28-Bam-R (Table S2). The amplified DNA fragment was digested with NdeI and BamHI (Takara) and cloned into pET-28b digested with the same restriction enzymes using the Ligation Mix (Nippon Gene, Tokyo, Japan). The resulting plasmid was named pET28-His-CsgA (Table S1).

The DNA fragment containing *csgA* without its stop codon was amplified from the genomic DNA of BW25113 using PCR with KOD Plus Neo DNA polymerase and the primer set CsgA-His-pET28-F and CsgA-His-pET28-R (Table S2). Plasmid pET-28b was linearized using inverse PCR with KOD Plus Neo DNA polymerase and the primer set Inverse-pET28-F and Inverse-pET28-R (Table S2). Amplified DNA fragments were ligated using the GeneArt Seamless Cloning Kit (Thermo Fisher Scientific, Waltham, MA, USA). The resulting plasmid was named pET28-CsgA-His (Table S1).

The mutation plasmid pET28-His-CsgA^A130G/S131G^ was constructed using inverse PCR-based site-specific mutagenesis with KOD Plus Neo DNA polymerase and the primer set CsgA-A130G/S131G-F and CsgA-A130G/S131G-R (Table S2).

DNA fragments encoding C-terminally His_6_-tagged DegP (DegP-His), C-terminally His_6_- tagged DegQ (DegQ-His), C-terminally His_6_-tagged Prc (Prc-His), and non-tagged YdgD were amplified from the genomic DNA of BW25113 by PCR using KOD Plus Neo DNA polymerase and the primer sets DegP-His-pCA24N-F1 and DegP-His-pCA24N-R1, DegQ-His-pCA24N-F1 and DegQ-His-pCA24N-R1, Prc-His-pCA24N-F1 and Prc-His-pCA24N-R1, and YdgD-pCA24N-F and YdgD-pCA24N-R, respectively (Table S2). The amplified DNA fragments were cloned into pCA24N using GeneArt Seamless Cloning Kit. The resulting plasmids were named pDegP-His, pDegQ-His, pPrc-His, and pYdgD, respectively (Table S1).

The mutation plasmids pDegP-S210A-His, pDegQ-S187A-His, and pPrc-S452A-His (Table S1), which express the protease activity-deficient mutants DegP^S210A^-His, DegQ^S187A^-His, and Prc^S452A^-His, were constructed using inverse PCR-based site-specific mutagenesis with KOD Plus Neo DNA polymerase and the primer sets DegP-S210A-F and DegP-S210A-R, DegQ-S187A-F and DegQ-S187A-R, and Prc-S452A-F and Prc-S452A-R, respectively (Table S2).

The DNA fragments encoding CsgA-sfGFP and sfGFP were amplified from pBAD- CsgBA-sfGFP (*13*) using PCR with KOD Plus Neo DNA polymerase and the primer sets CsgA-sfGFP-pCold-F and CsgA-sfGFP-pCold-R and sfGFP-pCold-F and sfGFP-pCold-R, respectively (Table S2). The DNA fragment encoding mCherry was amplified from pBAD-CsgB-mCherry/CsgA-sfGFP (*13*) using PCR with the mCherry-pCold-F and mCherry-pCold-R primer sets (Table S2). The DNA fragments encoding CpxP and YjfN were amplified from the genome of BW25113 using PCR with KOD Plus Neo DNA polymerase and the primer sets CpxP-pCold-F and CpxP-pCold-R and YjfN-pCold-F and YjfN-pCold-R, respectively (Table S2). The amplified DNA fragments were cloned into linearized pCold I using the GeneArt Seamless Cloning Kit. The resulting plasmids were named pCold-CsgA-sfGFP, pCold-sfGFP, pCold-mCherry, pCold-CpxP, and pCold-YjfN, respectively (Table S1).

The primers used in this study are listed in Table S2. The sequences of the constructed plasmids were confirmed using DNA sequencing (Eurofins Genomics, Tokyo, Japan).

### Antibodies

Anti-CsgA (generated by MBL, Tokyo, Japan), anti-CsgG (generated by MBL), anti-GFP (MBL), anti-MBP (Abcam, Cambridge, MA, USA), anti-OmpA (gifted by Dr. Yoshinori Akiyama), anti-penta-His (Qiagen), anti-PNPase (generated by Scrum, Tokyo, Japan), anti-RFP (MBL), and anti-RpoD (Abcam) antibodies were used as previously reported (*13, 14*). Horseradish peroxidase (HRP)-conjugated goat anti-rabbit and anti-mouse IgG (BioRad Laboratories) secondary antibodies were used. Rabbit polyclonal anti-DegP, anti-DegQ, and anti-Prc antibodies were developed by Eurofins Genomics, using purified proteins as antigens.

### Protein purification

Unmodified CsgA was purified from extracellular curli fibers, as previously reported (Chapman, et al. 2002), with some modifications. BW25113 was cultured on YESCA agar plates (1% casamino acid, 0.1% yeast extract, 2% agar) at 26 ± 2°C for 72 h. Colonies were harvested using a scraper and suspended in 10 mM Tris-HCl (pH 8.0). The cells were homogenized on ice using a homogenizer (1 min, 1 min interval). After centrifugation at 5,000 ×g for 10 min at 23–25°C, bacterial cells were removed. The supernatant was incubated with 150 mM NaCl for 30 min on ice and centrifuged at 15,000×g for 10 min at 4°C. The pellet was washed three times with 10 mM Tris-HCl (pH 8.0) and 150 mM NaCl, and the purified curli was suspended in 10 mM Tris-HCl (pH 8.0) and stored at −80°C. Prior to performing the protein degradation assay, the frozen curli sample was thawed on ice and centrifuged at 15,000 ×g for 10 min at 4°C. The resulting pellet was resuspended in 100–200 μL of HFIP and incubated at 23–25°C for 10 min. Following incubation, HFIP was removed by vacuum centrifugation, and the remaining pellet was resuspended in an 8 M urea solution. The purity and concentration of CsgA were verified by SDS-PAGE. Just before initiating the protein degradation assay, urea was removed using a ZetaSpin column (Thermo Fisher Scientific) at 4°C, and the eluted CsgA was used as unfolded CsgA for the assay.

N-terminally His_6_-tagged CsgA (His-CsgA), N-terminally His_6_-tagged CsgA^A130G/S131G^ (His-CsgA^A130G/S131G^), and C-terminally His_6_-tagged CsgA (CsgA-His) were purified under denaturing conditions. *E. coli* LOBSTR cells harboring pET28-CsgA-His, pET28- His-CsgA, or pET28-His-CsgA^A130G/S131G^ were cultured at 37°C in 1 L of LB medium supplemented with 50 μg/mL Km. When the optical density at 600 nm wavelength reached 0.5–0.6, the culture was supplemented with 1 mM IPTG and further incubated at 37°C for 1 h to induce these proteins. Bacterial cells were harvested by centrifugation at 8,000 ×g for 10 min at 4°C, suspended in lysis denaturation buffer [10 mM Tris-HCl (pH 8.0) and 6 M guanidine-hydrochloride (Gdn-HCl)], and disrupted by stirring overnight at 23–25°C. After centrifugation at 20,000 ×g for 30 min at 4°C, the supernatant was passed through a 0.45 μm filter. The cleared lysate was incubated with Ni-NTA resin (1 mL bed volume) for at least 1 h at 4°C. The bound proteins were washed with 50 mL of the lysis denaturation buffer and eluted with 10 mL of the elution-denaturation buffer [20 mM Tris-HCl (pH 8.0), 250 mM imidazole, 6 M Gdn-HCl]. Similar to untagged CsgA described above, Gdn-HCl was removed using a ZetaSpin column (Thermo Fisher Scientific) at 4°C just before performing the protein degradation assay, and the eluted His-CsgA and CsgA-His were used as unfolded substrates for the assay.

C-terminally His_6_-tagged DegP (DegP-His), DegP (DegQ-His), Prc (Prc-His), and Prc^S452A^(Prc^S452A^-His) were purified as follows: BL21 (DE3) cells harboring pDegP-His, pDegQ- His, pPrc-His, or pPrc-S452A-His were cultured at 37°C in 2 L of LB medium supplemented with 30 μg/mL chloramphenicol (Cm). When optical density at 600 nm reached 0.5–0.6, the culture was supplemented with 1 mM IPTG and further incubated at 37°C for 3 h to induce these proteins. Bacterial cells were harvested by centrifugation at 5,000 ×g for 10 min at 4°C, suspended in lysis buffer [10 mM Tris-HCl (pH 8.0), 150 mM NaCl, 10 mM imidazole], and disrupted by sonication. After centrifugation at 20,000 ×g for 30 min at 4°C, the supernatant was passed through a 0.45 μm filter. The cleared lysate was incubated with TALON resin (1 mL bed volume) for at least 1 h at 4°C. The bound proteins were washed with 50 mL of the lysis buffer and eluted with 10 mL of the elution buffer [20 mM Tris-HCl (pH 8.0), 150 mM NaCl, 250 mM imidazole]. After buffer exchange with buffer A [20 mM Tris-HCl (pH 8.0), 1 mM DTT, 10% glycerol] using PD- 10 column, the proteins were purified by anion-exchange chromatography using a Mono Q column with linear gradient of 0–500 mM NaCl. Fractions containing the proteins of interest were pooled and concentrated using an Amicon Ultra ultrafiltration device (30, 50, or 100 K). If purity was not sufficient, proteins were further purified by size exclusion chromatography using Superdex 75 or Superdex 200 size exclusion columns in buffer B [20 mM Tris-HCl (pH 8.0), 300 mM NaCl, 1 mM DTT, 10% glycerol].

N-terminal His_6_-tagged CsgA-sfGFP (His-CsgA-sfGFP), sfGFP (His-sfGFP), mCherry (His-mCherry), CpxR (His-CpxP), and YjfN (His-YjfN) were purified as follows: *E. coli* BL21(DE3) cells harboring pCold-CsgA-sfGFP, pCold-sfGFP, pCold-mCherry, pCold-CpxP, or pCold-YjfN were grown at 37°C in 2 L of LB medium supplemented with 100 μg/mL Amp. When optical density at 600 nm reached 0.5–0.6, the culture was incubated at 15°C for 30 min, and 1 mM IPTG was added. After further incubation at 15°C overnight, the bacterial cells were harvested by centrifugation at 5,000 ×g for 10 min at 4°C. The pellets were resuspended in lysis buffer and disrupted by sonication. After centrifugation at 20,000 ×g for 30 min at 4°C, the supernatant was passed through a 0.45 μm filter. The cleared lysate was incubated with TALON resin (1 mL bed volume) for at least 1 h at 4°C. The bound proteins were washed with 50 mL of the lysis buffer and eluted with 10 mL of the elution buffer. After buffer exchange with buffer A using a PD- 10 column, the proteins were purified by anion-exchange chromatography using a Mono Q column with a linear gradient of 0–500 mM NaCl. Fractions containing the proteins of interest were pooled and concentrated using an Amicon Ultra ultrafiltration device (10, 30, or 50 K). If the purity was insufficient, the proteins were further purified using size exclusion chromatography with Superdex 75 or Superdex 200 size exclusion columns in buffer B.

Purity of proteins were confirmed using 15% SDS-PAGE. The protein concentration was determined using 660 nm Protein Assay Reagent (Invitrogen).

### Curli production

*E. coli* strains were cultured in LB medium at 30°C overnight. The overnight cultures were spotted on CR-containing YESCA agar [1% casamino acids, 0.1% yeast extract, 2% agar, 10 μg/mL CR, 1 μg/mL Coomassie Brilliant Blue (CBB)], and the plates were incubated at 26 ± 2°C for 3 days. Colony images were captured using a digital camera or scanner (EPSON, Tokyo, Japan). When required, the colonies were collected and subjected to immunoblotting.

### Multi-copy suppressor screening

*E. coli* BW25113 Δ*degP* Δ*degQ* was transformed with the ASKA clone plasmids expressing one of the potential periplasmic proteases (BepA, DacA, DacB, DacC, DacD, IAP, MepA, NlpC, PbpG, Prc, PtrA, TesA, UmuD, YafL, YdcP, YdgD, YdhO, YebA, YhbU, and YhjJ). The transformants were grown at 26 ± 2°C for 3 days on CR-YESCA plates supplemented with 30 μg/mL Cm and 0.01–1 mM IPTG. White colonies overexpressing a periplasmic protease responsible for CsgA degradation in the periplasm were selected as candidates. Colony PCR was performed using GoTaq DNA polymerase (Promega, Madison, Wisconsin, USA) to amplify the corresponding protease genes. The amplified DNA was analyzed using agarose gel electrophoresis and sequenced (Eurofins Genomics).

### Cell fractionation

Cell fractionation was conducted based on the protocol by Malherbe et al. (*36*), with modifications. *E. coli* cells were grown on YESCA plates at 28°C for 48 h and collected in test tubes. The bacterial cells were weighed, and 50 mg of the cells were suspended in 500 μL of ice-cold PBS supplemented with 100 U/mL DNase I (Roche Diagnostics, Mannheim, Germany) to digest extracellular DNA in the colonies. The mixture was vortexed, and a 100 μL aliquot was taken as the total fraction. The remaining suspension (400 μL) was centrifuged at 20,000 ×g for 3 min at 4°C. The supernatant was collected into a new test tube as the extracellular fraction.

The bacterial pellet was resuspended in the same volume (400 μL) of spheroplast buffer [100 mM Tris-HCl (pH 8.0), 500 mM (17.11%) sucrose, 0.5 mM EDTA] and incubated for 5 min at room temperature. After centrifugation at 20,000 ×g for 3 min at 4°C, the supernatant was discarded, and the pellet was resuspended in 400 μL of 1 mM MgCl_2_ and incubated for 15 sec on ice. After centrifugation at 10,000 ×g for 3 min at 4°C, the supernatant was collected as the periplasmic fraction. The pellet was washed once with 400 μL of 1 mM MgCl_2_.

The cells were resuspended in 400 μL of cytoplasmic buffer [10 mM Tris-HCl (pH 8.0), 2.5 mM EDTA] and disrupted by sonication (five times for 10 sec each) on ice. Unbroken cells and debris were removed by centrifugation at 2,000 ×g for 5 min at 4°C. The supernatant was further centrifuged at 10,000 ×g for 10 min at 4°C. The insoluble fraction was used as the aggregate fraction. The soluble fraction, containing the cytoplasm and membrane, was ultracentrifuged at 120,000 ×g for 45 min at 4°C to separate the soluble cytoplasmic fraction and the insoluble membrane fraction. The membrane fraction was dissolved in 400 μL of cytoplasmic buffer by brief sonication.

The aggregate fraction was processed according to Tomoyasu et al. (*37*). The aggregate fraction was resuspended in 320 μL of cytoplasmic buffer by brief sonication; then, 80 μL of 10% (v/v) NP-40 was added. The aggregated proteins were isolated by centrifugation (15,000 ×g for 15 min at 4°C). This washing procedure was repeated to remove contaminating membrane proteins. NP-40-insoluble pellets were washed with 400 μL of cytoplasmic buffer and resuspended in the same volume by brief sonication.

Equivalent volumes of the total, membrane, periplasm, cytoplasm, and aggregate fractions were separated by SDS-PAGE on SDS-15% polyacrylamide gels. CsgA and other marker proteins were detected by immunoblotting. The fluorescence of sfGFP in the separated fractions was measured using an Infinite F200 Pro (Tecan, Männedorf, Switzerland) microtiter plate reader.

### Immunoblotting

Immunoblotting was performed to detect curli-related proteins, as previously reported (*13, 14*). Macrocolonies formed on YESCA or CR-YESCA plates were collected and their weights were measured. Bacterial cell suspensions were prepared in PBS at a concentration of 0.1 mg/μL. Ten microliters of the suspension was diluted in 70 μL of 2 × SDS-PAGE sample buffer (Fujifilm Wako, Osaka, Japan) and boiled at 95°C for 5 min. The samples were subjected to SDS-PAGE with 15% SDS polyacrylamide gels. The proteins were transferred from the gel to a PVDF membrane using an iBlot 2 dry blotting system (Thermo Fisher Scientific) following the manufacturer’s instructions. The membranes were blocked in 0.1% Tween 20-containing Tris-buffered saline (TBS-T) (10 mM Tris-HCl (pH7.4), 100 mM NaCl) supplemented with 1% skimmed milk for 30 min at 23–25°C or overnight at 4°C and then incubated with primary antibodies in CanGet Signal solution 1 (Toyobo, Osaka, Japan) for 30 min at 23–25°C or overnight at 4°C. After washing twice with TBS-T for 5 min at 23–25°C, the membranes were incubated with secondary antibodies in CanGet Signal solution 2 (Toyobo) for 30 min at 23–25°C or overnight at 4°C. After washing three times with TBS-T for 5 min at 23–25°C, proteins were detected using ECL Prime (Cytiva) and LAS-4000 image analyzer (Cytiva). Anti-CsgA (1:1,000), anti-CsgG (1:1,000), anti-DegP (1:10,000), anti-DegQ (1:10,000), anti-GFP (1:1,000), anti-MBP (1:5,000), anti-OmpA (1:10,000), anti-penta-His (1:2,000), anti-Prc (1:10,000), anti-PNPase (1:10,000), anti-RFP (1:1,000), and anti-RpoD (1:2,000) were used as primary antibodies. HRP-conjugated goat anti-rabbit IgG (1:50,000) and HRP-conjugated goat anti-mouse IgG (1:10,000) were used as secondary antibodies.

### Quantitative real-time-polymerase reaction (qRT-PCR)

To measure transcripts of *csgA* and *csgD*, pRT-PCR was performed as previously described (*13, 14, 38, 39*).*E. coli* BW25113 and its isogenic mutant cells were grown overnight in LB medium at 30°C with shaking. Aliquots (30 μL) of the cultures were spotted on YESCA plate and incubated at 26 ± 2°C for 24 h. Total RNA was purified using the QIAGEN RNeasy Mini Kit (Qiagen). cDNA was synthesized using the PrimeScript II 1st Strand cDNA Synthesis Kit (Takara, Otsu, Japan). Transcript levels of *csgA* and *csgD* were measured by qRT-PCR using primers RT-csgA-F/RT-csgA-R and RT-csgD-F/RT-csgD-R, respectively (Table S2). As an internal control, *ftsZ* transcript levels were measured using the primers RT-ftsZ-F and RT-ftsZ-R (Table S2). qRT-PCR was performed using TB Green Premix Ex Taq™ II (Tli RNase H Plus) (Takara) and Step One Plus (Applied Biosystems, Foster City, CA, USA).

### Protease and peptidase assays

Unfolded CsgA-His (2 μM) and His-CsgA (2 μM) were incubated with purified Prc-His (2 μM), Prc^S542A^-His (2 μM), DegP-His (2 μM), or DegQ-His (2 μM) in buffer C [20 mM HEPES-KOH (pH 7.6), 100 mM KCl, 20 mM MgCl_2_, and 1 mM EDTA] at 37°C for 60 min. Unfolded CsgA (5 μM) was incubated with Prc-His (5 μM), Prc^S542A^-His (5 μM), DegP-His (5 μM), or DegQ-His (5 μM) in buffer C at 37°C for 30 min. His-CsgA^A130G/S131G^ (5 μM) was incubated with Prc-His (5 μM) in buffer C at 37°C for 30 min. At the indicated time points, the reaction was stopped by adding an equal volume of 2 × SDS-PAGE sample buffer. The samples were applied to 15% SDS polyacrylamide gels. The gels were then stained with CBB.

To analyze the effects of adaptor and activator proteins on degradation of CsgA by DegP and DegQ, His-CsgA (5 μM), and HFIP-depolymerized CsgA (5 μM) were incubated with DegP-His (5 μM) or DegQ-His (5 μM) in the presence and absence of His-YjfN (25 μM) or His-CpxP (25 μM) in buffer C at 37°C for the indicated times. At the indicated time points, the reaction was stopped by adding an equal volume of 2 × SDS-PAGE sample buffer. CsgA degradation was analyzed using SDS-PAGE followed by CBB staining. Degradation of His-CsgA was further analyzed using immunoblotting with anti-His and anti-CsgA antibodies as mentioned above.

### GFP-Trap assay

CsgA-binding proteins were isolated and identified using GFP-Trap magnetic agarose (ChromoTek, Munich, Germany). BA-FP Δ*degP* Δ*degQ* was cultured on YESCA agar plates at 26 ± 2°C for 2 days. As a control, BW25113 harboring pBAD-CsgA_1-20_-sfGFP (*13*) was used. After harvesting cells using a scraper, bacterial cells were suspended in lysis buffer [10 mM Tris-Cl (pH 7.5), 150 mM NaCl, 0.5 mM EDTA, 0.5% NP-40, protease inhibitor cocktail (Nacalai Tesque, Kyoto, Japan)] and disrupted by sonication. The broken bacterial cells were centrifuged at 20,000 ×g for 10 min at 4°C, and the resulting supernatants containing sfGFP and CsgA_21-131_-sfGFP were incubated with GFP-Trap magnetic agarose for 1 h at 4°C. After washing the agarose with washing buffer [10 mM Tris-Cl (pH 7.5), 150 mM NaCl, 0.5 mM EDTA, 0.05% NP-40, protease inhibitor cocktail], bound proteins were eluted with 2 × SDS-PAGE sample buffer. The eluted fractions were analyzed using SDS-PAGE, followed by LC-MS/MS analysis.

### LC-MS/MS analysis

To identify CsgA-binding proteins, LC-MS/MS analysis was performed to identify proteins co-purified with CsgA_21-131_-sfGFP and sfGFP using MS Bioworks LLC (Ann Arbor, MI, USA).

For determination of Prc cleavage sites in CsgA, CsgA (50 μM) was incubated with Prc (5 μM) in buffer C at 37°C for 0, 10, 30, and 60 min. At these time points, the reaction was stopped by mixing equal volumes of 2 × SDS-PAGE sample buffer. The degradation products were analyzed using SDS-PAGE and LC-MS/MS. The CBB-stained Prc protein and degradation product bands were excised from the SDS-PAGE gel, cut into approximately 1-1.5 mm sized cubes for efficient in-gel trypsin digestion, and de-stained. The proteins in the gel pieces were reduced using DTT (Thermo Fisher Scientific), alkylated with iodoacetamide (Thermo Fisher Scientific), and digested with trypsin and lysyl-endopeptidase (Promega). The resultant peptides were analyzed on an Advance UHPLC system (Michrom Bioresources, Auburn, California, USA) coupled to a Q Exactive mass spectrometer (Thermo Fisher Scientific) with the raw data processed using Xcalibur (Thermo Fisher Scientific). Raw data were analyzed against the SwissProt or UniProt KB databases restricted to *E. coli* using Proteome Discoverer version 1.4 (Thermo Fisher Scientific) with the Mascot search engine version 2.7 (Matrix Science, London, UK). A decoy database comprising either randomized or reversed sequences in the target database was used to estimate the false discovery rate (FDR) estimation, and the percolator algorithm was used to evaluate false positives. Search results were filtered against 1% global FDR for high confidence level. Peptide sequences were also determined from MS/MS spectra by PEAKS 8.5 (Bioinformatics Solutions, Waterloo, Canada; parent mass error tolerance 10.0 ppm, fragment mass error tolerance, 0.02 Da) and manually checked.

After incubating CsgA (50 μM) with Prc (5 μM) at 37°C for 0, 10, 30, and 60 min, each incubated mixture was directly analyzed on an Advance UHPLC system (Michrom Bioresources, Auburn, California, USA) coupled to a Q Exactive mass spectrometer (Thermo Fisher Scientific) with the raw data processed using Xcalibur (Thermo Fisher Scientific). Two microliters of 100-fold diluted sample were trapped on a pre-column (L-column 2 ODS, 5 µm, 0.3 × 5 mm, CERI) then de-salted and concentrated. The mixture was separated on micro-ODS column (L-column 2 ODS, 3 µm, 0.1 × 150 mm, CERI), at 40°C with flow rate 500 nL/min. Mobile phase A consisted of 0.1% formic acid, and mobile phase B comprised ACN containing 0.1% formic acid. The eluent gradient ranged from 5% to 65% B over 0–60 min and then increased from 65% to 95% B from 60 to 70 min. MS spectra were acquired at m/z 350–1,500, with a resolution of 70,000. The resolution of the MS/MS spectra was 17,500 using Top-5 data-dependent acquisition. The values of high-energy collision dissociation were set at 20, 25, and 30 normalized collisional energy).

### Fluorescent microscopy

Fluorescent microscopy was performed as previously reported (Sugimoto, et al. 2018). Briefly, overnight cultures (50–100 μL) of the BA-FP and its derivative strains were spotted on YESCA plates containing 2% agar, and the plates were incubated at 26 ± 2°C for 3 days. Macro colonies were suspended in sterilized PBS (-), and the suspensions (5 μL) were placed on slide glasses and covered with coverslips. The fluorescence signals of sfGFP and mCherry in these samples were visualized using a fluorescence microscope (Nikon, Tokyo, Japan) equipped with B2 (excitation filter: 450–490 nm; barrier filter: 520 nm) and G2A (excitation filter: 510–560 nm; barrier filter: 590 nm) filters. Phase-contrast images were also captured.

### Statistical analysis

Statistical analyses were performed using the GraphPad Prism Ver. 9 for Windows OS (GraphPad Software, Inc., San Diego, CA, USA). One-way ANOVA was used to determine whether the groups exhibited statistically significant differences. Multiple comparisons with a control group were performed using one-way ANOVA with Tukey’s correction (Fig. 1, D and F). For the comparison of more than eight groups, one-way ANOVA with two-stage linear step-up procedure of Benjamini, Krieger, and Yekutieli correction was performed with a False Discovery Rate of 5% (Fig. 5, A and B). All experiments were performed at least three times to ensure the accuracy and reproducibility of the results. *P* < 0.05 was considered to indicate statistical significance for all analyses.

## Acknowledgments

We acknowledge Mrs. Naoko Toda for her exceptional experimental support. We also extend our thanks to Dr. Yoshinori Akiyama (Kyoto University) for providing the anti-OmpA antisera and to the National BioResource Project of Japan for supplying the KEIO collection mutants and the ASKA clone library.

## Funding

JSPS Grant-in-Aid for Fund for the Promotion of Joint International Research [Fostering Joint International Research (A)] 18KK0429 (SS)

JST FOREST Program JPMJFR23719171 (SS)

JST ERATO JPMJER1502 (SS)

Kumamoto University, IMEG, the Joint Usage/Research Center for Developmental Medicine (SS)

Kumamoto University, IMEG, the program of the Inter-University Research Network for High Depth Omics (SS, NT, KY)

The Uehara Memorial Foundation (SS)

The Jikei University Strategic Prioritizing Research Fund (SS, YK)

## Author contributions

Conceptualization: SS, YK

Methodology: SS, NT

Investigation: SS, YT, NT

Visualization: SS Supervision: KY, YK

Writing—original draft: SS

Writing—review & editing: SS, YT, NT, YK, KY

## Data and materials availability

All data are available in the main text or the supplementary materials.

## Supplementary Materials

Figs. S1 to S5,

Tables S1 to S2.

## References

1. G. G. Jayaraj, M. S. Hipp, F. U. Hartl, Functional Modules of the Proteostasis Network. Cold Spring Harb Perspect Biol 12, (2020).

2. D. Balchin, M. Hayer-Hartl, F. U. Hartl, In vivo aspects of protein folding and quality control. Science 353, aac4354 (2016).

3. N. Jain, M. R. Chapman, Bacterial functional amyloids: Order from disorder. Biochim Biophys Acta Proteins Proteom 1867, 954–960 (2019).

4. M. R. Chapman et al., Role of Escherichia coli curli operons in directing amyloid fiber formation. Science 295, 851–855 (2002).

5. B. Cao et al., Structure of the nonameric bacterial amyloid secretion channel. Proc Natl Acad Sci U S A 111, E5439–5444 (2014).

6. P. Goyal et al., Structural and mechanistic insights into the bacterial amyloid secretion channel CsgG. Nature 516, 250–253 (2014).

7. L. S. Robinson, E. M. Ashman, S. J. Hultgren, M. R. Chapman, Secretion of curli fibre subunits is mediated by the outer membrane-localized CsgG protein. Mol Microbiol 59, 870–881 (2006).

8. M. Zhang, H. Shi, X. Zhang, X. Zhang, Y. Huang, Cryo-EM structure of the nonameric CsgG-CsgF complex and its implications for controlling curli biogenesis in Enterobacteriaceae. PLoS Biol 18, e3000748 (2020).

9. N. D. Hammer, J. C. Schmidt, M. R. Chapman, The curli nucleator protein, CsgB, contains an amyloidogenic domain that directs CsgA polymerization. Proc Natl Acad Sci U S A 104, 12494–12499 (2007).

10. A. A. Nenninger, L. S. Robinson, S. J. Hultgren, Localized and efficient curli nucleation requires the chaperone-like amyloid assembly protein CsgF. Proc Natl Acad Sci U S A 106, 900–905 (2009).

11. H. M. Swasthi, J. L. Basalla, C. E. Dudley, A. G. Vecchiarelli, M. R. Chapman, Cell surface-localized CsgF condensate is a gatekeeper in bacterial curli subunit secretion. Nat Commun 14, 2392 (2023).

12. E. A. Epstein, M. A. Reizian, M. R. Chapman, Spatial clustering of the curlin secretion lipoprotein requires curli fiber assembly. J Bacteriol 191, 608–615 (2009).

13. S. Sugimoto et al., Multitasking of Hsp70 chaperone in the biogenesis of bacterial functional amyloids. Commun Biol 1, 52 (2018).

14. S. Sugimoto et al., Hierarchical Model for the Role of J-Domain Proteins in Distinct Cellular Functions. J Mol Biol 433, 166750 (2021).

15. M. L. Evans et al., The bacterial curli system possesses a potent and selective inhibitor of amyloid formation. Mol Cell 57, 445–455 (2015).

16. A. A. Nenninger et al., CsgE is a curli secretion specificity factor that prevents amyloid fibre aggregation. Mol Microbiol 81, 486–499 (2011).

17. M. L. Evans et al., E. coli chaperones DnaK, Hsp33 and Spy inhibit bacterial functional amyloid assembly. Prion 5, 323–334 (2011).

18. H. Loferer, M. Hammar, S. Normark, Availability of the fibre subunit CsgA and the nucleator protein CsgB during assembly of fibronectin-binding curli is limited by the intracellular concentration of the novel lipoprotein CsgG. Mol Microbiol 26, 11–23 (1997).

19. S. Grau et al., Implications of the serine protease HtrA1 in amyloid precursor protein processing. Proc Natl Acad Sci U S A 102, 6021–6026 (2005).

20. T. Clausen, M. Kaiser, R. Huber, M. Ehrmann, HTRA proteases: regulated proteolysis in protein quality control. Nat Rev Mol Cell Biol 12, 152–162 (2011).

21. M. Kitagawa et al., Complete set of ORF clones of Escherichia coli ASKA library (a complete set of E. coli K-12 ORF archive): unique resources for biological research. DNA Res 12, 291–299 (2005).

22. S. K. Singh, S. Parveen, L. SaiSree, M. Reddy, Regulated proteolysis of a cross-link-specific peptidoglycan hydrolase contributes to bacterial morphogenesis. Proc Natl Acad Sci U S A 112, 10956–10961 (2015).

23. D. D. Isaac, J. S. Pinkner, S. J. Hultgren, T. J. Silhavy, The extracytoplasmic adaptor protein CpxP is degraded with substrate by DegP. Proc Natl Acad Sci U S A 102, 17775–17779 (2005).

24. S. Kim, I. Song, G. Eom, S. Kim, A Small Periplasmic Protein with a Hydrophobic C-Terminal Residue Enhances DegP Proteolysis as a Suicide Activator. J Bacteriol 200, (2018).

25. H. Hara, Y. Yamamoto, A. Higashitani, H. Suzuki, Y. Nishimura, Cloning, mapping, and characterization of the Escherichia coli prc gene, which is involved in C-terminal processing of penicillin-binding protein 3. J Bacteriol 173, 4799–4813 (1991).

26. K. C. Keiler, P. R. Waller, R. T. Sauer, Role of a peptide tagging system in degradation of proteins synthesized from damaged messenger RNA. Science 271, 990–993 (1996).

27. K. R. Silber, K. C. Keiler, R. T. Sauer, Tsp: a tail-specific protease that selectively degrades proteins with nonpolar C termini. Proc Natl Acad Sci U S A 89, 295–299 (1992).

28. G. Jubelin et al., CpxR/OmpR interplay regulates curli gene expression in response to osmolarity in Escherichia coli. J Bacteriol 187, 2038–2049 (2005).

29. A. Vianney et al., Escherichia coli tol and rcs genes participate in the complex network affecting curli synthesis. Microbiology (Reading) 151, 2487–2497 (2005).

30. D. L. Hung, T. L. Raivio, C. H. Jones, T. J. Silhavy, S. J. Hultgren, Cpx signaling pathway monitors biogenesis and affects assembly and expression of P pili. EMBO J 20, 1508–1518 (2001).

31. C. G. Evans, S. Wisen, J. E. Gestwicki, Heat shock proteins 70 and 90 inhibit early stages of amyloid beta-(1-42) aggregation in vitro. J Biol Chem 281, 33182–33191 (2006).

32. S. Edvardson et al., A deleterious mutation in DNAJC6 encoding the neuronal-specific clathrin-uncoating co-chaperone auxilin, is associated with juvenile parkinsonism. PLoS One 7, e36458 (2012).

33. S. Poepsel et al., Determinants of amyloid fibril degradation by the PDZ protease HTRA1. Nat Chem Biol 11, 862–869 (2015).

34. K. A. Datsenko, B. L. Wanner, One-step inactivation of chromosomal genes in Escherichia coli K-12 using PCR products. Proc Natl Acad Sci U S A 97, 6640–6645 (2000).

35. T. Baba et al., Construction of Escherichia coli K-12 in-frame, single-gene knockout mutants: the Keio collection. Mol Syst Biol 2, 2006 0008 (2006).

36. G. Malherbe, D. P. Humphreys, E. Dave, A robust fractionation method for protein subcellular localization studies in Escherichia coli. Biotechniques 66, 171–178 (2019).

37. T. Tomoyasu, A. Mogk, H. Langen, P. Goloubinoff, B. Bukau, Genetic dissection of the roles of chaperones and proteases in protein folding and degradation in the Escherichia coli cytosol. Mol Microbiol 40, 397–413 (2001).

38. K. Arita-Morioka, K. Yamanaka, Y. Mizunoe, T. Ogura, S. Sugimoto, Novel strategy for biofilm inhibition by using small molecules targeting molecular chaperone DnaK. Antimicrob Agents Chemother 59, 633–641 (2015).

39. K. I. Arita-Morioka et al., Inhibitory effects of Myricetin derivatives on curli-dependent biofilm formation in Escherichia coli. Sci Rep 8, 8452 (2018).

