## Supplementary Materials for "Periplasmic serine protease Prc is responsible for amyloid subunit CsgA degradation and proteostasis in *Escherichia coli*"

Shinya Sugimoto *et al.*

**This PDF file includes:**

Figs. S1 to S5
Tables S1 to S2

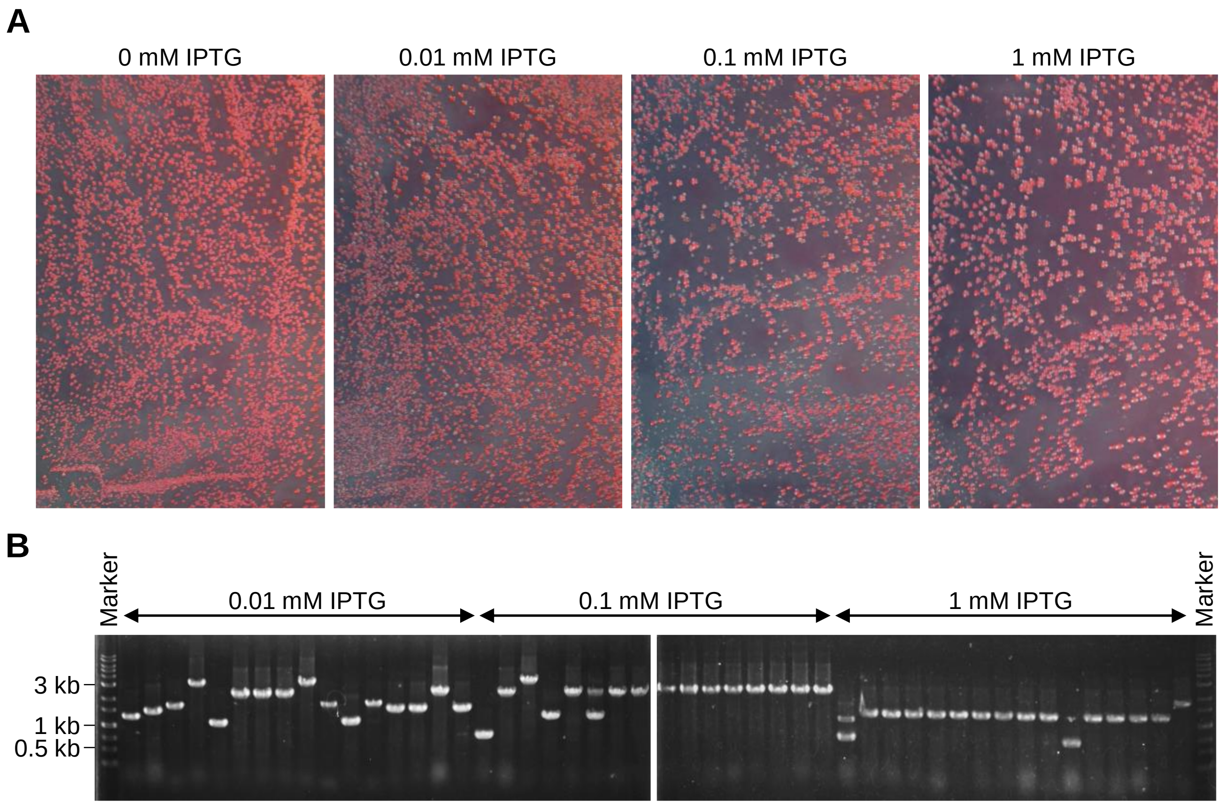


**Fig. S1. Identification of periplasmic proteases responsible for CsgA degradation in the periplasm.** (**A**) Pictures of *E. coli* colonies grown on Congo Red-containing YESCA plates supplemented with 30 μg/mL chloramphenicol and the indicated concentrations of IPTG for 3 days at 26 ± 2°C. BW25113 Δ*degP* Δ*degQ* was transformed with the ASKA clone plasmids expressing one of the potential periplasmic proteases (BepA, DacA, DacB, DacC, DacD, IAP, MepA, NlpC, PbpG, Prc, PtrA, TesA, UmuD, YafL, YdcP, YdgD, YdhO, YebA, YhbU, and YhjJ). White colonies were selected to perform colony PCR and DNA sequencing for identification of the genes encoded on the plasmids. (**B**) Amplified DNA fragments by colony PCR were analyzed using agarose gel electrophoresis.

**A**

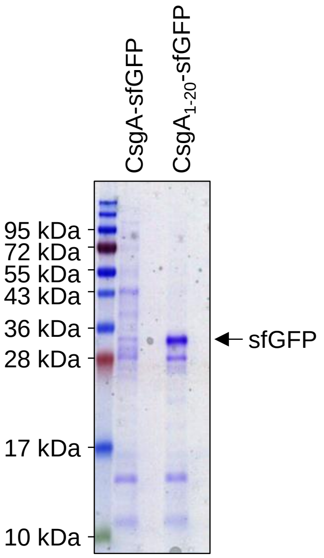


**B**

| Identified Proteins | Localization | MW | IP | Control | Fold Change (IP / Control) |
|---|---|---|---|---|---|
| Curli production assembly/transport component CsgG | OM | 31 kDa | 113 | 0 | NA |
| Uncharacterized protease YdcP | PP | 73 kDa | 36 | 0 | NA |
| Probable protease SohB | IM | 39 kDa | 23 | 0 | NA |
| Signal peptidase I (LepB) | IM | 36 kDa | 16 | 0 | NA |
| Beta-barrel assembly-enhancing protease (BepA) | PP | 54 kDa | 14 | 0 | NA |
| Curli production assembly/transport component CsgF | OM | 15 kDa | 12 | 0 | NA |
| Probable L,D-transpeptidase YnhG | ND | 36 kDa | 11 | 0 | NA |
| Probable L,D-transpeptidase YcbB | ND | 68 kDa | 6 | 0 | NA |
| Probable L,D-transpeptidase YcfS | ND | 35 kDa | 6 | 0 | NA |
| Protease 4 (SspA) | IM | 67 kDa | 5 | 0 | NA |
| Curli assembly protein CsgC | PP | 12 kDa | 5 | 0 | NA |
| Major curli subunit CsgA | EX | 15 kDa | 175 | 3 | 58.3 |
| ATP-dependent zinc metalloprotease FtsH | IM | 71 kDa | 101 | 2 | 50.5 |
| Minor curli subunit CsgB | EX | 16 kDa | 41 | 2 | 20.5 |
| Modulator of FtsH protease HflC | IM | 38 kDa | 49 | 5 | 9.8 |
| Probable L,D-transpeptidase ErfK/SrfK | ND | 34 kDa | 45 | 5 | 9 |
| Modulator of FtsH protease HflK | IM | 46 kDa | 69 | 11 | 6.27 |
| Cytosol aminopeptidase | CP | 55 kDa | 31 | 6 | 5.17 |
| Metalloprotease TldD | CP | 51 kDa | 9 | 2 | 4.5 |
| ATP-dependent Clp protease ATP-binding subunit ClpX | CP | 46 kDa | 43 | 11 | 3.91 |
| Lon protease | CP | 87 kDa | 199 | 61 | 3.26 |
| Tail-specific protease Prc | PP | 77 kDa | 19 | 6 | 3.17 |
| ATP-dependent Clp protease ATP-binding subunit ClpA | CP | 84 kDa | 157 | 50 | 3.14 |
| Gamma-glutamyltranspeptidase | ND | 62 kDa | 12 | 4 | 3 |

**Fig. S2. Identification of CsgA-binding proteins using GFP-Trap assay.** (**A**) GFP-trap assay was performed to co-purify CsgA-sfGFP and its biding proteins using anti-GFP nanobody-conjugated magnetic beads. Isolated proteins from BW25113 Δ*degP* Δ*degQ* expressing CsgA-sfGFP were analyzed using SDS-PAGE. BW25113 Δ*degP* Δ*degQ* expressing C-terminally sfGFP-fused signal sequence of CsgA (CsgA$_{1\text{-}20}$-sfGFP) was used as a negative control. CsgA$_{1\text{-}20}$-sfGFP is translocated to the periplasm, and the secretion signal is removed. Proteins were identified using LC-MS/MS analysis. (**B**) Identified CsgA-binding proteins are listed. CsgA and known CsgA-binding proteins are colored in blue. Periplasmic proteases are colored in red. IP, the numbers of the identified peptides of proteins co-purified with CsgA-sfGFP. Control, the numbers of the identified peptides of proteins co-purified with sfGFP. Fold change > 3 was considered as specific biding. CP, cytoplasm; PP, periplasm; IM, inner membrane; OM, outer membrane; EX, extracellular; NA, not applicable; ND, not determined.

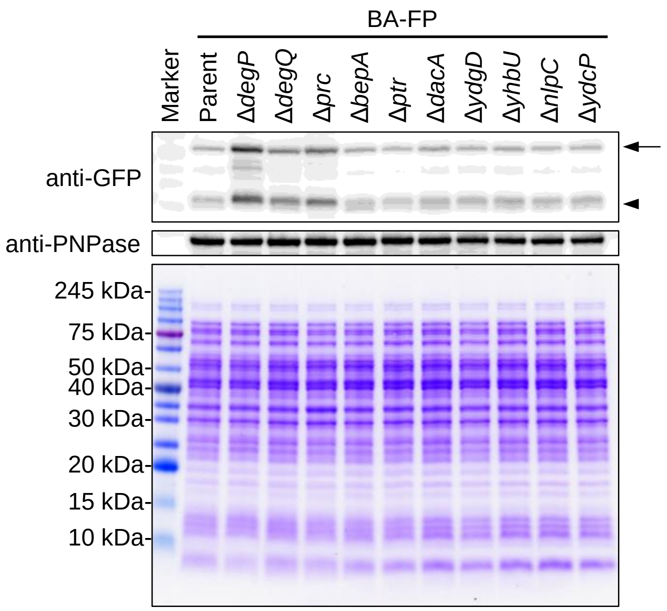


**Fig. S3. Immunoblotting to detect CsgA-sfGFP in the BA-FP derivative strains.** CsgA-sfGFP in the indicated single-gene deletion mutans was detected using immunoblotting with anti-GFP antibody. PNPase was detected as a loading control. The black arrows and arrowheads represent full-length CsgA-sfGFP and sfGFP, respectively. The positions of molecular mass markers are indicated at the left.

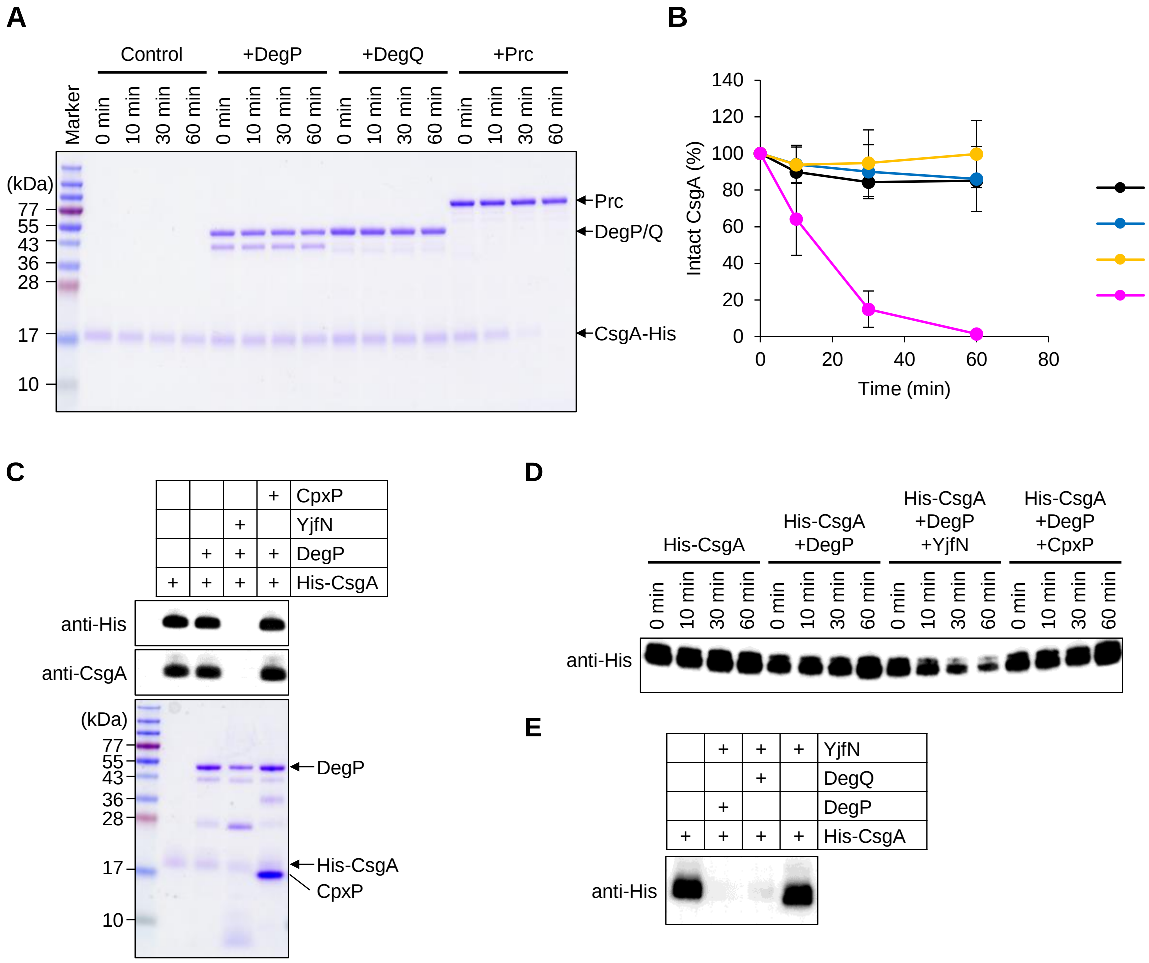


**Fig. S4. DegP and DegQ degrade CsgA in a suicide activator-dependent mechanism.** (**A**) In vitro degradation of purified C-terminally His-tagged CsgA (CsgA-His) by purified DegP, DegQ, and Prc analyzed using SDS-PAGE. (**B**) The band intensity of CsgA was quantified using the ImageQuant TL software. The intensity at 0 min were set as 100%. Data are represented as mean ± SD from three independent experiments. (**C**) Degradation of N-terminally His-tagged CsgA (His-CsgA) by DegP in the presence and absence of suicide activator YjfN and adaptor CpxP. After 24 h incubation, the samples were analyzed using SDS-PAGE and immunoblotting using anti-His and anti-CsgA antibodies. (**D**) Time-dependent degradation of His-CsgA in the presence of the indicated proteins analyzed using immunoblotting with anti-His antibody. (**E**) Degradation of CsgA by DegP and DegQ in the presence of YjfN. After 24 h incubation, the samples were analyzed using immunoblotting with anti-His antibody.

**21-151**:
GVVPQYGGGGNHGGGGNNSGPNSELNIYQYGGGNSALALQTDARNSDLTITQHGGGNGADVGQGSDDSSIDLTQRGFGNS
ATLDQWNGKNSEMTVKQFGGGNGAAVDQTASNSSVNVTQVGFGNNATAHQY
$C_{544}H_{833}N_{173}O_{204}S_1$

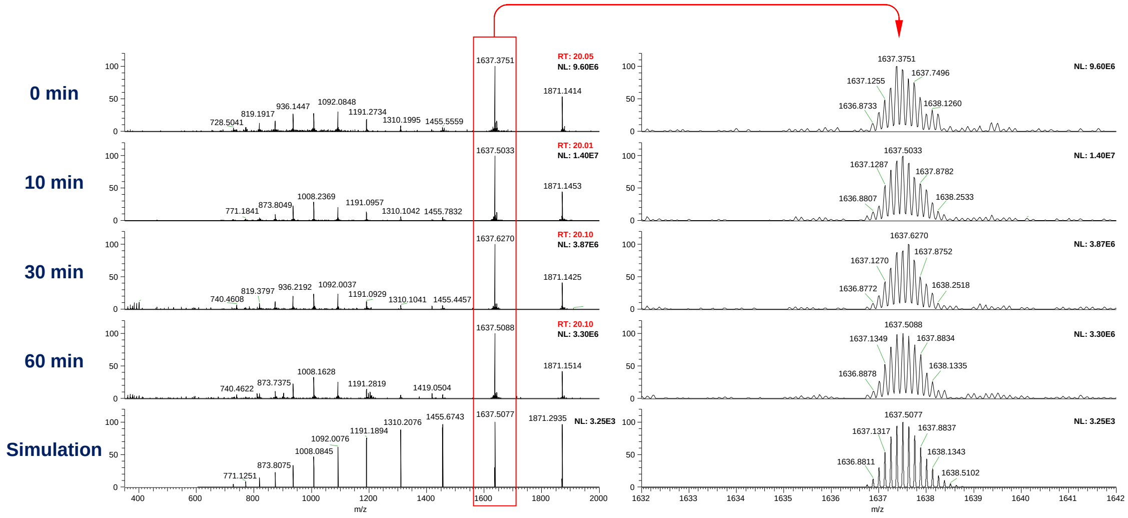


**21-93**:
GVVPQYGGGGNHGGGGNNSGPNSELNIYQYGGGNSALALQTDARNSDLTITQHGGGNGADVGQGSDDSSIDLT
$C_{291}H_{449}N_{93}O_{115}$

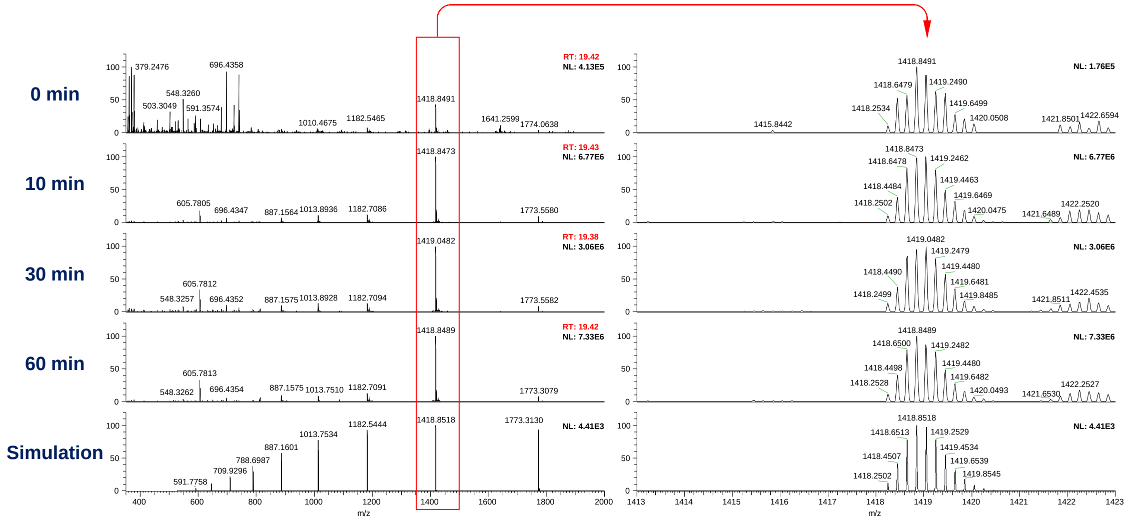

**21-58**:
GVVPQYGGGGNHGGGGNNSGPNSELNIYQYGGGNSALA
**$C_{151}H_{224}N_{48}O_{55}$**

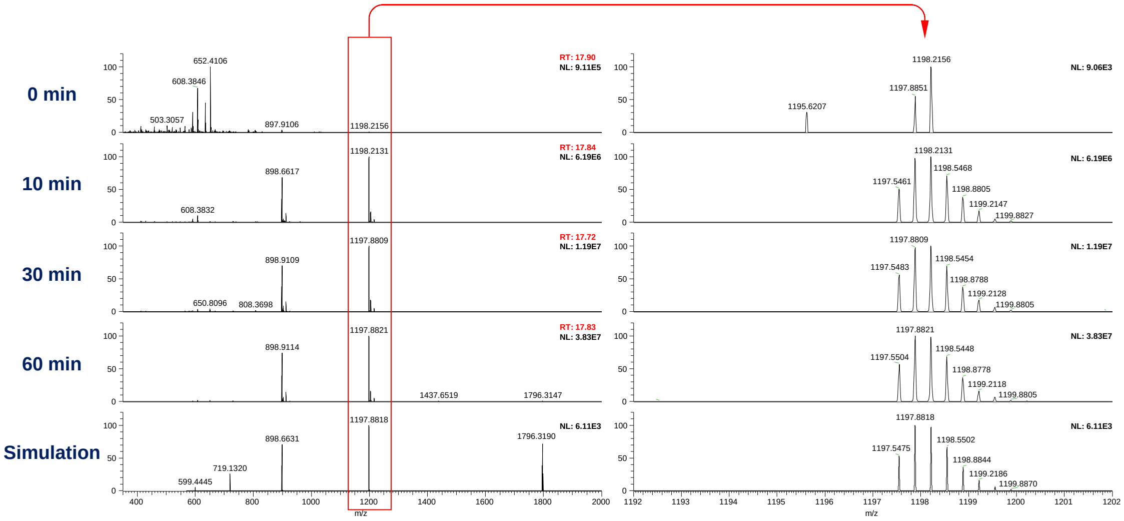


**59-71**:
LQTDARNSDLTIT
**$C_{59}H_{102}N_{18}O_{24}$**

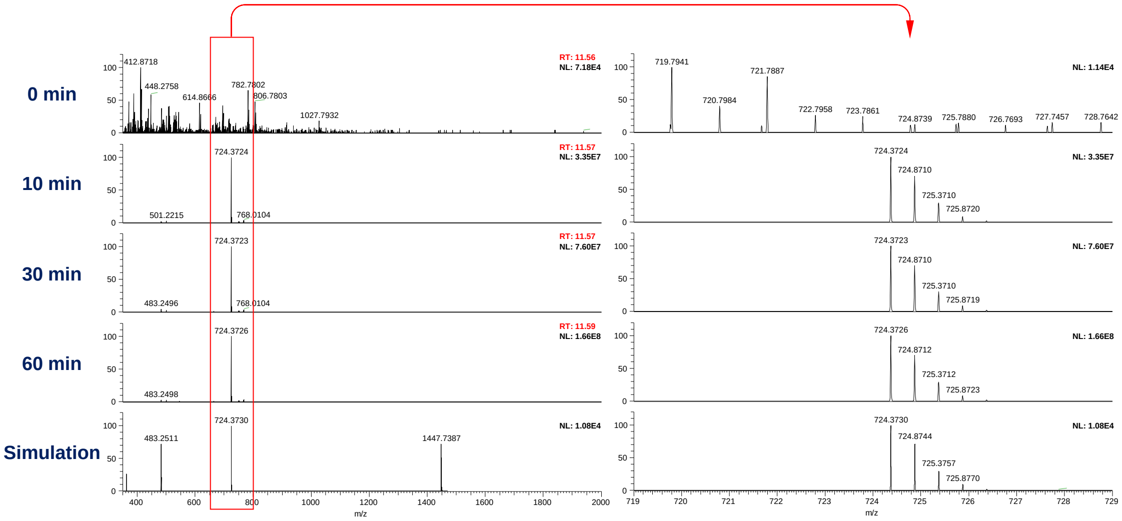

**72-93**:
QHGGGNGADVGQGSDDSSIDLT
**$C_{81}H_{127}N_{27}O_{38}$**

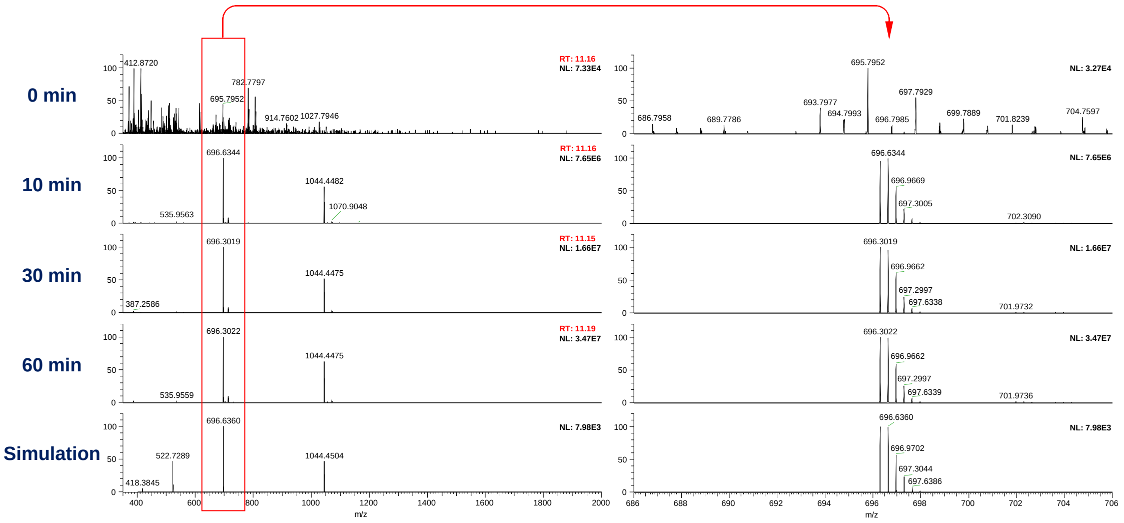


**94-111**:
QRGFGNSATLDQWNGKNS
**$C_{83}H_{126}N_{28}O_{29}$**

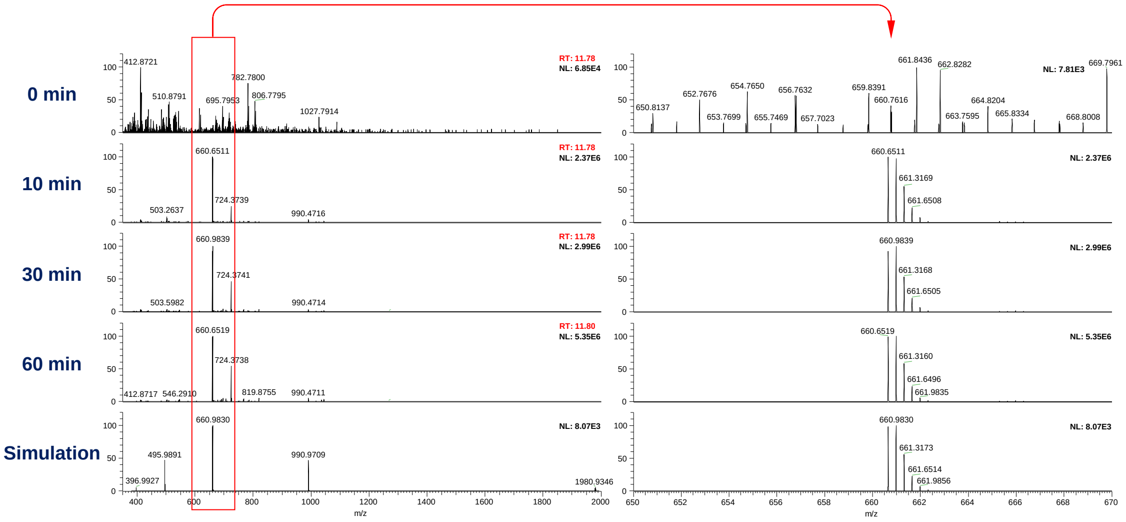

**94-114**:
QRGFGNSATLDQWNGKNSEMT
**$C_{97}H_{149}N_{31}O_{35}S_1$**

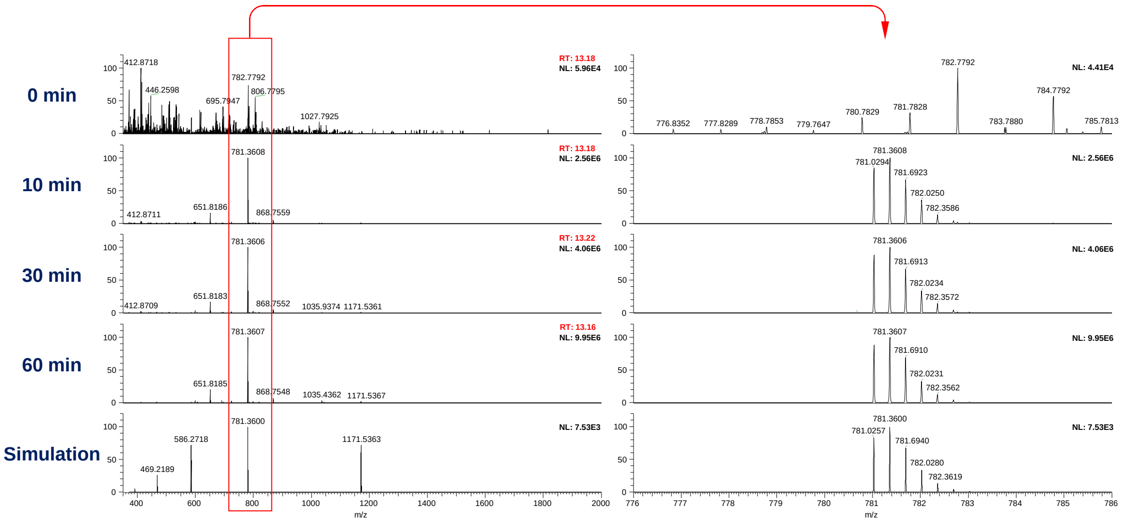

**94-115**:
QRGFGNSATLDQWNGKNSEMTV
**$C_{102}H_{158}N_{32}O_{36}S_1$**

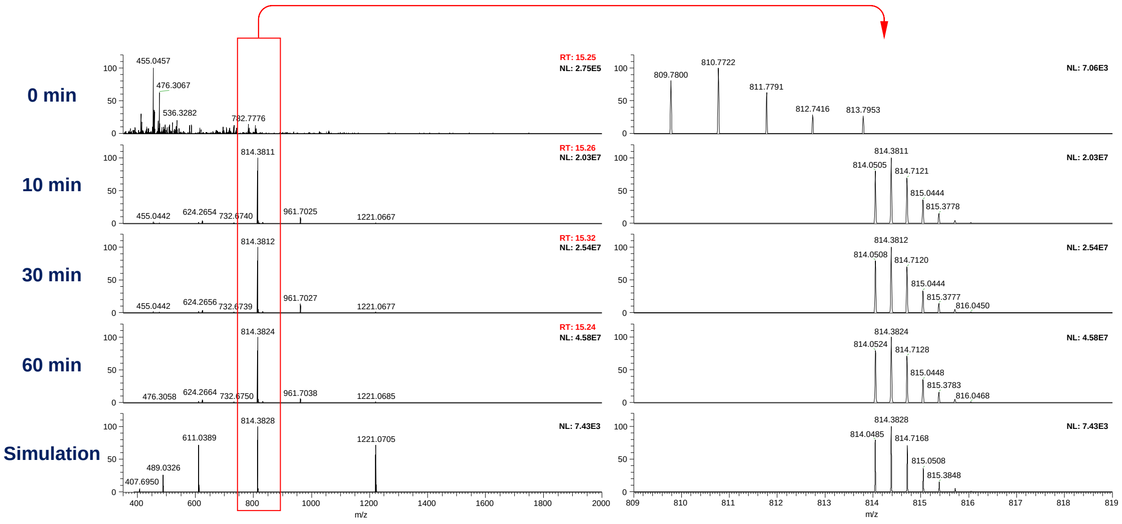

**94-130**:
QRGFGNSATLDQWNGKNSEMTVKQFGGGNGAAVDQTA
$\mathbf{C_{161}H_{249}N_{51}O_{57}S_1}$

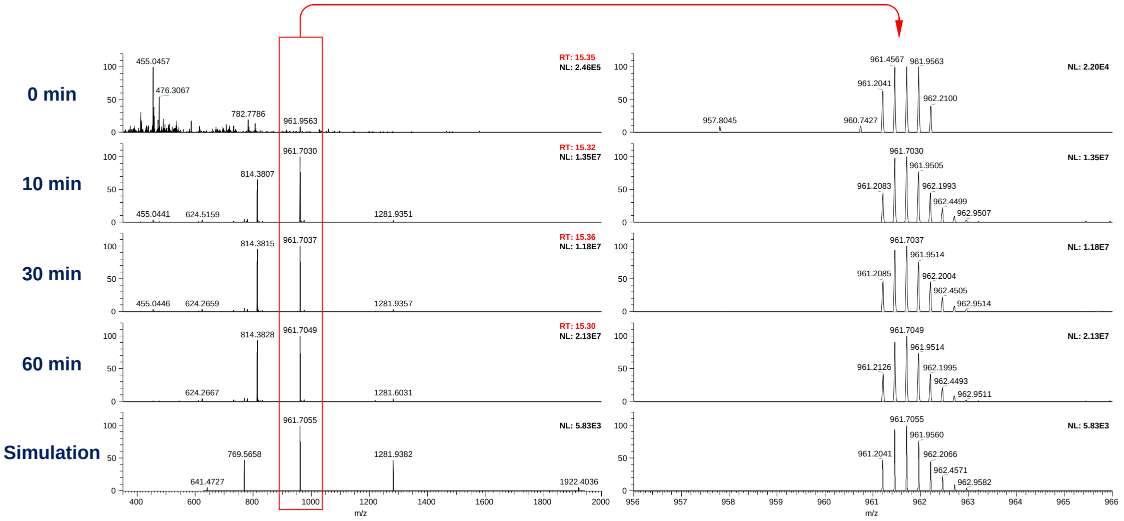


**94-151**:
QRGFGNSATLDQWNGKNSEMTVKQFGGGNGAAVDQTASNSSVNVTQVGFGNNATAHQY
$\mathbf{C_{253}H_{386}N_{80}O_{90}S_1}$

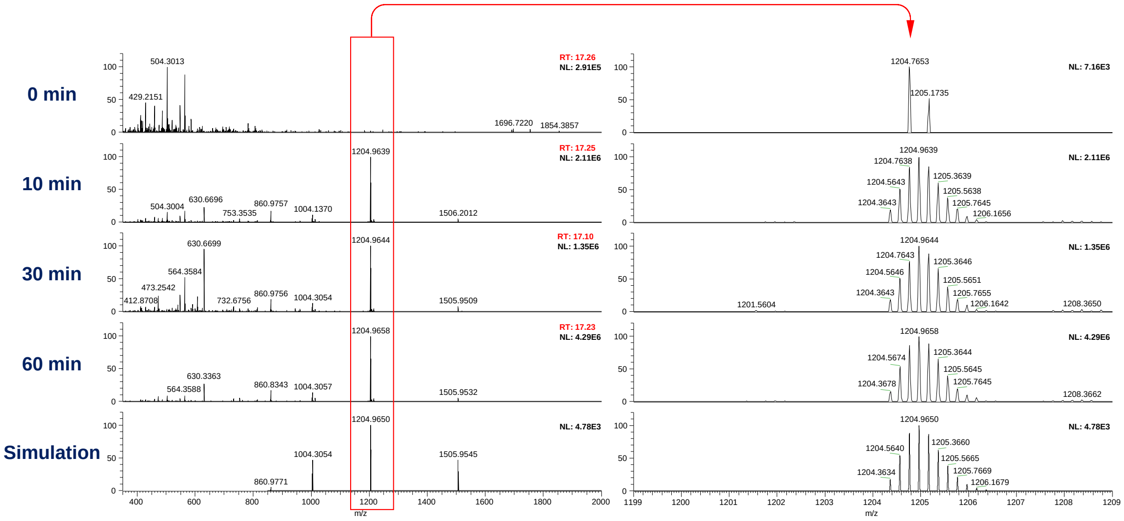

**102-115**:
TLDQWNGKNSEMTV
$\mathbf{C_{68}H_{107}N_{19}O_{25}S_1}$

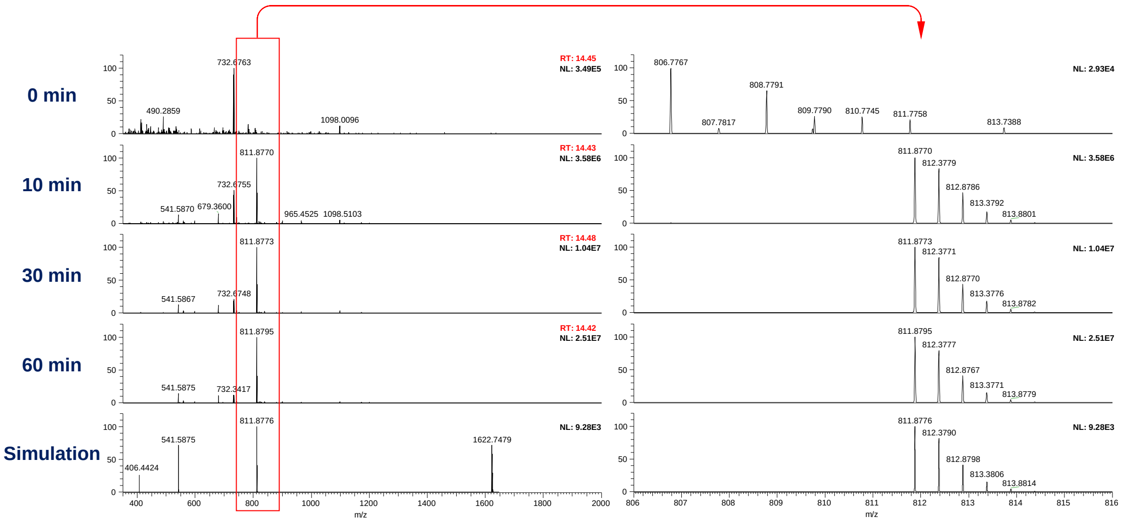


**115-130**:
VKQFGGGNGAAVDQTA
$\mathbf{C_{64}H_{102}N_{20}O_{23}}$

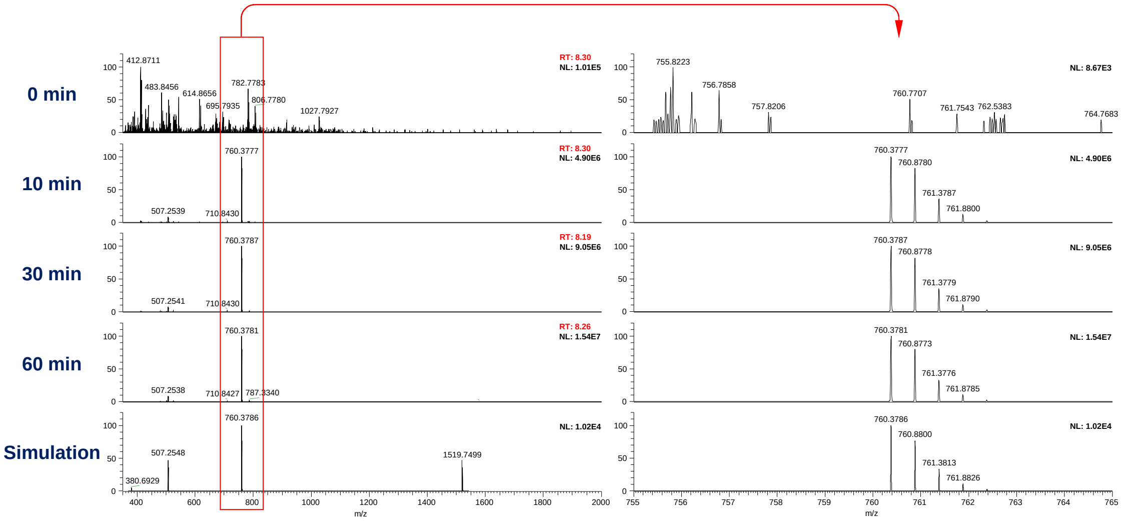

**116-130**:
KQFGGGNGAAVDQTA
$\mathbf{C_{59}H_{93}N_{19}O_{22}}$

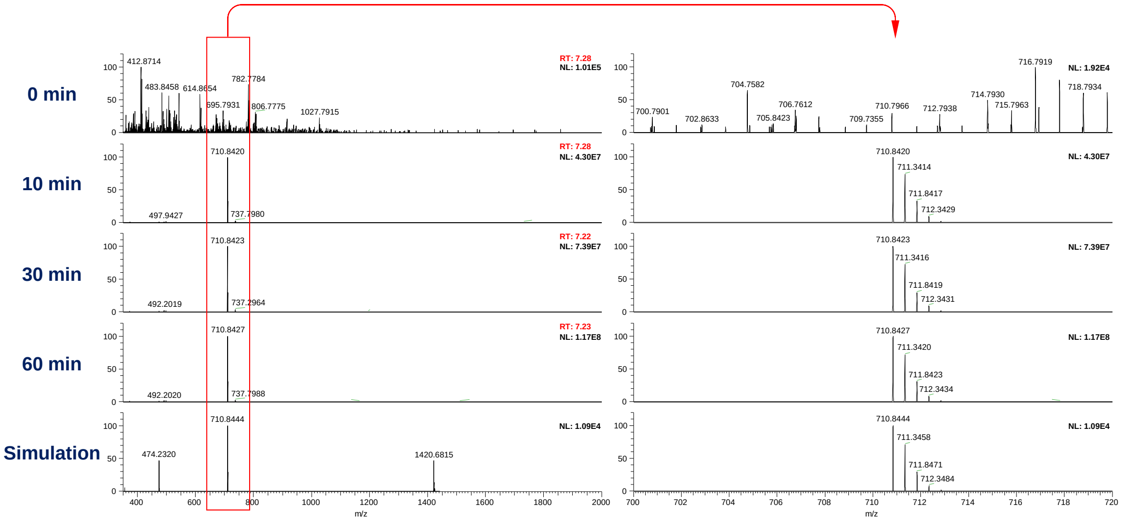


**131-151**:
SNSSVNVTQVGFGNNATAHQY
$\mathbf{C_{92}H_{139}N_{29}O_{34}}$

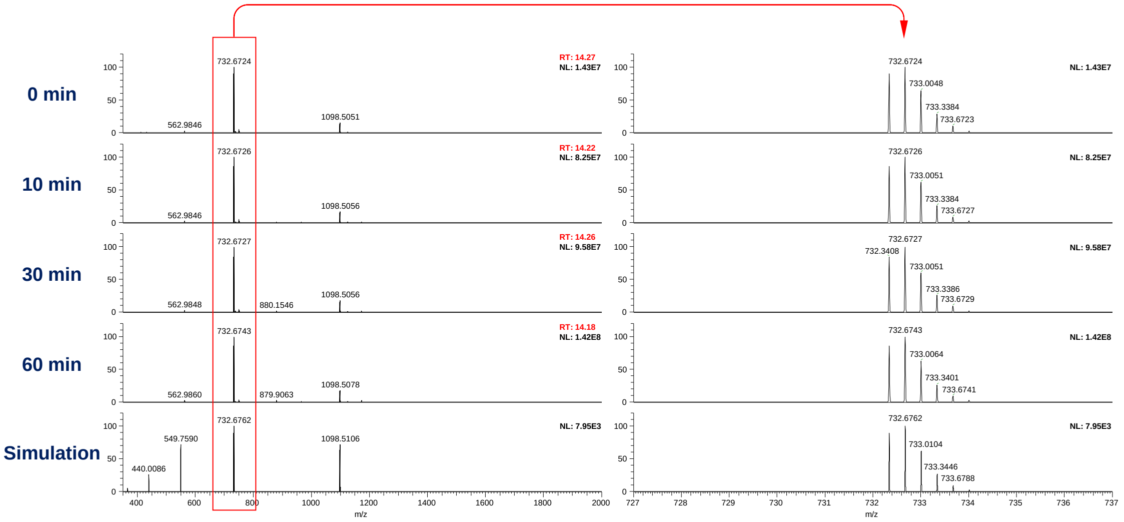

**139-151**:
QVGFGNNATAHQY
**$C_{61}H_{87}N_{19}O_{20}$**

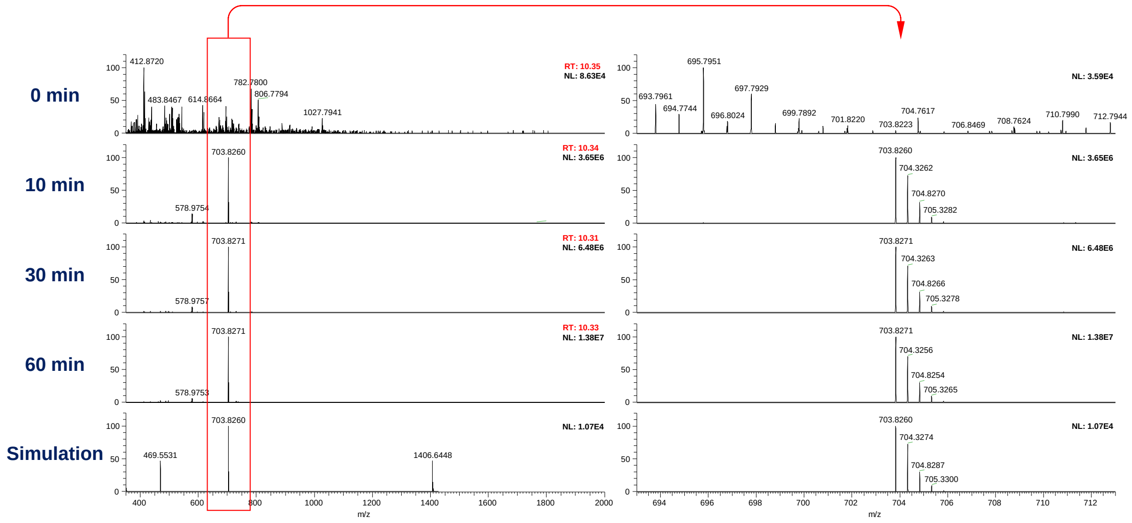


**Fig. S5. Mass spectra of Prc protein and degradation product obtained by incubating CsgA with Prc.** Mass spectra were extracted from each peak of MS chromatograms in Fig. 3F, and main multi-charged ion peak was expanded to observe isotopic distribution spectrum. Mass spectra of simulation were obtained from creating a simulated isotopic distribution spectrum of a chemical formulas of CsgA and degradation product using Xcalibur Qual Browser (Thermo Fisher Scientific).

**Table S1.**
Bacterial strains and plasmids used in this study.

| **Strains and plasmids** | **Description** | **Source/Reference** |
|---|---|---|
| ***E. coli* strains** | | |
| DH5α | *F-, Φ80dlacZΔM15, Δ(lacZYA-argF)U169, deoR, recA1, endA1, hsdR17(rK-, mK+), phoA, supE44, λ-, thi-1, gyrA96, relA1* | Toyobo |
| BL21(DE3) | $F^-$ *ompT hsdS*$_B$(*r*$_B^-$ *m*$_B^-$) *gal(λcI857 ind1 Sam7 nin5 lacUV5-T7gene1) dcm* (DE3) | Novagen |
| LOBSTR | *E. coli* expression strain derived from BL21(DE3) for elimination of common contaminants from His-tag purification | Kerafast |
| BW25113 | K-12 wild type strain of Keio collection | Baba et al. 2006 (*35*) |
| JW0157-KC | BW25113 Δ*degP*::Km$^R$ | Baba et al. 2006 (*35*) |
| JW0627-KC | BW25113 Δ*dacA*::Km$^R$ | Baba et al. 2006 (*35*) |
| JW1020-KC | BW25113 Δ*csgG*::Km$^R$ | Baba et al. 2006 (*35*) |
| JW1021-KC | BW25113 Δ*csgF*::Km$^R$ | Baba et al. 2006 (*35*) |
| JW1022-KC | BW25113 Δ*csgE*::Km$^R$ | Baba et al. 2006 (*35*) |
| JW1023-KC | BW25113 Δ*csgD*::Km$^R$ | Baba et al. 2006 (*35*) |
| JW1024-KC | BW25113 Δ*csgB*::Km$^R$ | Baba et al. 2006 (*35*) |
| JW1025-KC | BW25113 Δ*csgA*::Km$^R$ | Baba et al. 2006 (*35*) |
| JW1026-KC | BW25113 Δ*csgC*::Km$^R$ | Baba et al. 2006 (*35*) |
| JW1431-KC | BW25113 Δ*ydcP*::Km$^R$ | Baba et al. 2006 (*35*) |
| JW1590-KC | BW25113 Δ*ydgD*::Km$^R$ | Baba et al. 2006 (*35*) |
| JW1698-KC | BW25113 Δ*nlpC*::Km$^R$ | Baba et al. 2006 (*35*) |
| JW1732-KC | BW25113 Δ*spy*::Km$^R$ | Baba et al. 2006 (*35*) |
| JW2205-KC | BW25113 Δ*rcsB*::Km$^R$ | Baba et al. 2006 (*35*) |
| JW2479-KC | BW25113 Δ*bepA*::Km$^R$ | Baba et al. 2006 (*35*) |
| JW2789-KC | BW25113 Δ*ptr*::Km$^R$ | Baba et al. 2006 (*35*) |
| JW3132-KC | BW25113 Δ*nlpI*::Km$^R$ | Baba et al. 2006 (*35*) |
| JW3127-KC | BW25113 Δ*yhbU*::Km$^R$ | Baba et al. 2006 (*35*) |
| JW3203-KC | BW25113 Δ*degQ*::Km$^R$ | Baba et al. 2006 (*35*) |
| JW3883-KC | BW25113 Δ*cpxR*::Km$^R$ | Baba et al. 2006 (*35*) |
| Δ*prc* | BW25113 Δ*prc*::Km$^R$ | This study |
| Δ*degP* Δ*degQ* | BW25113 Δ*degP* Δ*degQ*::Km$^R$ | This study |
| Δ*degP* Δ*degQ* Δ*prc* | BW25113 Δ*degP* Δ*degQ* Δ*prc*::Km$^R$ | This study |
| Δ*csgG* Δ*degP* Δ*degQ* | BW25113 Δ*csgG* Δ*degP* Δ*degQ*::Km$^R$ | This study |
| Δ*csgG* Δ*prc* | BW25113 Δ*csgG* Δ*prc*::Km$^R$ | This study |
| Δ*csgG* Δ*degP* Δ*degQ* Δ*prc* | BW25113 Δ*csgG* Δ*degP* Δ*degQ* Δ*prc*::Km$^R$ | This study |
| Δ*degP* Δ*degQ* Δ*prc* Δ*rcsB* | BW25113 Δ*degP* Δ*degQ* Δ*prc* Δ*rcsB*::Km$^R$ | This study |
| Δ*degP* Δ*degQ* Δ*prc* Δ*cpxR* | BW25113 Δ*degP* Δ*degQ* Δ*prc* Δ*cpxR*::Km$^R$ | This study |
| Δ*degP* Δ*degQ* Δ*prc* Δ*rcsB* Δ*cpxR* | BW25113 Δ*degP* Δ*degQ* Δ*prc* Δ*rcsB* Δ*cpxR*::Km$^R$ | This study |
| Δ*rcsB* Δ*cpxR* | BW25113 Δ*rcsB* Δ*cpxR*::Km$^R$ | This study |
| Δ*csgG* Δ*csgC* | BW25113 Δ*csgG* Δ*csgC*:Km$^R$ | This study |
| Δ*csgG* Δ*prc* | BW25113 Δ*csgG* Δ*prc*::Km$^R$ | This study |
| Δ*csgG* Δ*prc* Δ*csgC* | BW25113 Δ*csgG* Δ*prc* Δ*csgC*::Km$^R$ | This study |
| Δ*csgG* Δ*rcsB* | BW25113 Δ*csgG* Δ*rcsB*::Km$^R$ | This study |

| | | |
|---|---|---|
| Δ*csgG* Δ*degP* Δ*degQ* Δ*rcsB* | BW25113 Δ*csgG* Δ*degP* Δ*degQ* Δ*rcsB*::Km$^R$ | This study |
| Δ*csgG* Δ*prc* Δ*rcsB* | BW25113 Δ*csgG* Δ*prc* Δ*rcsB*::Km$^R$ | This study |
| Δ*csgG* Δ*degP* Δ*degQ* Δ*prc* Δ*rcsB* | BW25113 Δ*csgG* Δ*degP* Δ*degQ* Δ*prc* Δ*rcsB*::Km$^R$ | This study |
| Δ*csgG* Δ*cpxR* | BW25113 Δ*csgG* Δ*cpxR*::Km$^R$ | This study |
| Δ*csgG* Δ*degP* Δ*degQ* Δ*prc* Δ*cpxR* | BW25113 Δ*csgG* Δ*degP* Δ*degQ* Δ*prc* Δ*cpxR*::Km$^R$ | This study |
| Δ*csgG* Δ*prc* Δ*cpxR* | BW25113 Δ*csgG* Δ*prc* Δ*cpxR*::Km$^R$ | This study |
| Δ*csgG* Δ*degP* Δ*degQ* Δ*prc* Δ*rcsB* Δ*cpxR* | BW25113 Δ*csgG* Δ*degP* Δ*degQ* Δ*prc* Δ*rcsB* Δ*cpxR*::Km$^R$ | This study |
| BA-FP | BW25113 *csgB-mcherry csgA-sfGFP* | This study |
| BA-FP Δ*degP* | BA-FP Δ*degP*::Km$^R$ | This study |
| BA-FP Δ*degQ* | BA-FP Δ*degQ*::Km$^R$ | This study |
| BA-FP Δ*degP degQ* | BA-FP Δ*degP degQ*::Km$^R$ | This study |
| BA-FP Δ*csgC* | BA-FP Δ*csgC*::Km$^R$ | This study |
| BA-FP Δ*spy* | BA-FP Δ*spy*::Km$^R$ | This study |
| BA-FP Δ*spy* Δ*csgC* | BA-FP Δ*spy* Δ*csgC*::Km$^R$ | This study |
| BA-FP Δ*spy* Δ*csgC* Δ*degP* Δ*degQ* | BA-FP Δ*spy* Δ*csgC* Δ*degP* Δ*degQ*::Km$^R$ | This study |
| BA-FP Δ*degP* Δ*degQ* Δ*bepA* | BA-FP Δ*degP* Δ*degQ* Δ*bepA*::Km$^R$ | This study |
| BA-FP Δ*degP* Δ*degQ* Δ*dacA* | BA-FP Δ*degP* Δ*degQ* Δ*dacA*::Km$^R$ | This study |
| BA-FP Δ*degP* Δ*degQ* Δ*nlpC* | BA-FP Δ*degP* Δ*degQ* Δ*nlpC*::Km$^R$ | This study |
| BA-FP Δ*degP* Δ*degQ* Δ*nlpI* | BA-FP Δ*degP* Δ*degQ* Δ*nlpI*::Km$^R$ | This study |
| BA-FP Δ*degP* Δ*degQ* Δ*prc* | BA-FP Δ*degP* Δ*degQ* Δ*prc*::Km$^R$ | This study |
| BA-FP Δ*degP* Δ*degQ* Δ*ptr* | BA-FP Δ*degP* Δ*degQ* Δ*ptr*::Km$^R$ | This study |
| BA-FP Δ*degP* Δ*degQ* Δ*ydcP* | BA-FP Δ*degP* Δ*degQ* Δ*ydcP*::Km$^R$ | This study |
| BA-FP Δ*degP* Δ*degQ* Δ*ydgD* | BA-FP Δ*degP* Δ*degQ* Δ*ydgD*::Km$^R$ | This study |
| BA-FP Δ*degP* Δ*degQ* Δ*yhbU* | BA-FP Δ*degP* Δ*degQ* Δ*yhbU*::Km$^R$ | This study |
| BA-FP Δ*bepA* | BA-FP Δ*bepA*::Km$^R$ | This study |
| BA-FP Δ*dacA* | BA-FP Δ*dacA*::Km$^R$ | This study |
| BA-FP Δ*nlpC* | BA-FP Δ*nlpC*::Km$^R$ | This study |
| BA-FP Δ*nlpI* | BA-FP Δ*nlpI*::Km$^R$ | This study |
| BA-FP Δ*prc* | BA-FP Δ*prc*::Km$^R$ | This study |
| BA-FP Δ*ptr* | BA-FP Δ*ptr*::Km$^R$ | This study |
| BA-FP Δ*ydcP* | BA-FP Δ*ydcP*::Km$^R$ | This study |
| BA-FP Δ*ydgD* | BA-FP Δ*ydgD*::Km$^R$ | This study |
| BA-FP Δ*yhbU* | BA-FP Δ*yhbU*::Km$^R$ | This study |
| **Plasmids** | | |
| pBAD/SS02 | Arabinose inducible vector, Amp$^R$ | Sugimoto et al. 2018 (*13*) |
| pCA24N | Empty vector of ASKA clone, Cm$^R$ | Kitagawa et al. 2005 (*21*) |
| pCold I | Cold shock expression vector with N-terminal His$_6$-tag, Amp$^R$ | Takara |
| pET-28b | pET vector with N-terminal His$_6$-tag, Km$^R$ | Novagen |
| pBAD-CsgBA-sfGFP | *csgBA* cloned in pBAD-sfGFP, Amp$^R$ | Sugimoto et al. 2018 (*13*) |
| pBAD-CsgBA$^{1–20}$-sfGFP | *csgB* and *csgA*$^{1–20}$ cloned in pBAD-sfGFP, Amp$^R$ | Sugimoto et al. 2018 (*13*) |
| pBAD-CsgB-mCherry/CsgA-sfGFP | *mcherry* cloned in pBAD-CsgBA-sfGFP, Amp$^R$ | Sugimoto et al. 2018 (*13*) |
| pCold-His-CsgA-sfGFP | *csgA-sfgfp* cloned in pCold I, Amp$^R$ | This study |
| pCold-His-CpxP | *cpxR* cloned in pCold I, Amp$^R$ | This study |
| pCold-His-mCherry | *mcherry* cloned in pCold I, Amp$^R$ | This study |

| | | |
|---|---|---|
| pCold-His-sfGFP | *sfgfp* cloned in pCold I, $Amp^R$ | This study |
| pCold-His-YjfN | *yjfN* cloned in pCold I, $Amp^R$ | This study |
| pCsgA-His | *csgA* cloned in pCA24N, C-terminal $His_6$-tag, $Cm^R$ | This study |
| pDegP-His | *degP* cloned in pCA24N, C-terminal $His_6$-tag, $Cm^R$ | This study |
| pDegP-S210A-His | A point mutation S210A introduced in pDegP-His, $Cm^R$ | This study |
| pDegQ-His | *degQ* cloned in pCA24N, C-terminal $His_6$-tag, $Cm^R$ | This study |
| pDegQ-S187A-His | A point mutation S210A introduced in pDegQ-His, $Cm^R$ | This study |
| pPrc-His | *prc* cloned in pCA24N, C-terminal $His_6$-tag, $Cm^R$ | This study |
| pPrc-S452A-His | A point mutation S210A introduced in pPrc-His, $Cm^R$ | This study |
| pYdgD | *ydgD* cloned in pCA24N, $Cm^R$ | This study |
| pBepA | *bepA* cloned in pCA24N, ASKA clone, $Cm^R$ | Kitagawa et al. 2005 (*21*) |
| pDacA | *dacA* cloned in pCA24N, ASKA clone, $Cm^R$ | Kitagawa et al. 2005 (*21*) |
| pDacB | *dacB* cloned in pCA24N, ASKA clone, $Cm^R$ | Kitagawa et al. 2005 (*21*) |
| pDacC | *dacC* cloned in pCA24N, ASKA clone, $Cm^R$ | Kitagawa et al. 2005 (*21*) |
| pDacD | *dacD* cloned in pCA24N, ASKA clone, $Cm^R$ | Kitagawa et al. 2005 (*21*) |
| pIAP | *iap* cloned in pCA24N, ASKA clone, $Cm^R$ | Kitagawa et al. 2005 (*21*) |
| pMepA | *mepA* cloned in pCA24N, ASKA clone, $Cm^R$ | Kitagawa et al. 2005 (*21*) |
| pNlpC | *nlpC* cloned in pCA24N, ASKA clone, $Cm^R$ | Kitagawa et al. 2005 (*21*) |
| pPbpG | *pbpG* cloned in pCA24N, ASKA clone, $Cm^R$ | Kitagawa et al. 2005 (*21*) |
| pPrc | *prc* cloned in pCA24N, ASKA clone, $Cm^R$ | Kitagawa et al. 2005 (*21*) |
| pPtr | *ptr* cloned in pCA24N, ASKA clone, $Cm^R$ | Kitagawa et al. 2005 (*21*) |
| pTesA | *tesA* cloned in pCA24N, ASKA clone, $Cm^R$ | Kitagawa et al. 2005 (*21*) |
| pUmuD | *umuD* cloned in pCA24N, ASKA clone, $Cm^R$ | Kitagawa et al. 2005 (*21*) |
| pYafL | *yafL* cloned in pCA24N, ASKA clone, $Cm^R$ | Kitagawa et al. 2005 (*21*) |
| pYdcP | *ydcP* cloned in pCA24N, ASKA clone, $Cm^R$ | Kitagawa et al. 2005 (*21*) |
| pYdhO | *ydhO* cloned in pCA24N, ASKA clone, $Cm^R$ | Kitagawa et al. 2005 (*21*) |
| pYebA | *yebA* cloned in pCA24N, ASKA clone, $Cm^R$ | Kitagawa et al. 2005 (*21*) |
| pYhbU | *yhbU* cloned in pCA24N, ASKA clone, $Cm^R$ | Kitagawa et al. 2005 (*21*) |
| pYhjJ | *yhjJ* cloned in pCA24N, ASKA clone, $Cm^R$ | Kitagawa et al. 2005 (*21*) |
| pET28-His-CsgA | *csgA* cloned in pET-28b, N-terminal $His_6$-tag, $Km^R$ | This study |
| pET28-His-$CsgA^{A130G/S131G}$ | *csgA*$^{A130G/S131G}$ cloned in pET-28b, N-terminal $His_6$-tag, $Km^R$ | This study |
| pET28-CsgA-His | *csgA* cloned in pET-28b, C-terminal $His_6$-tag, $Km^R$ | This study |

$Amp^R$, ampicillin resistance; $Cm^R$, chloramphenicol resistance; $Km^R$, kanamycin resistance.

**Table S2.**

Oligonucleotide primers used in this study.

| Name | Sequence (5' to 3') | Description | Reference |
|---|---|---|---|
| CsgB-mCherry-SOE-F | GCAGAGACAGTCGCAAATGGCTATTCGCGTGACACAACGT | Forward primer used for knockin of *csgB-mcherry* and *csgA-sfgfp* | This study |
| sfGFP-Km-SOE-R | GAAGCAGCTCCAGCCTACACTTATTTATACAGCTCATCCATACCATGCGTAATCCC | Reverse primer used for knockin of *csgB-mcherry* and *csgA-sfgfp* | This study |
| sfGFP-Km-SOE-F | GGGATTACGCATGGTATGGATGAGCTGTATAAATAAGTGTAGGCTGGAGCTGCTTC | Forward primer used for knockin of *csgB-mcherry* and *csgA-sfgfp* | This study |
| Km-CsgA-3'-SOE-R | AACAGGGCTTGCGCCCTGTTTCTGTAATACAAATGATGTACATATGAATATCCTCCTTA | Reverse primer used for knockin of *csgB-mcherry* and *csgA-sfgfp* | This study |
| CsgB-SOE-F | CAATTGTAGTGCAGAGACAGTCGCAAATGGCTATTCGCG | Forward primer used for knockin of *csgB-mcherry* and *csgA-sfgfp* | This study |
| CsgA-3'-R | CCCGAAAAAAAACAGGGCTTGCGCCCTGTTTCTGTAATAC | Reverse primer used for knockin of *csgB-mcherry* and *csgA-sfgfp* | This study |
| bepA-KO-F | GTGGGCTAATCTTCGCCGTTACACTCAAAG | Forward primer used for knockout of *bepA* | This study |
| bepA-KO-R | CGTTTTCTTTCAGCAGGTTCAGCGTTTCC | Reverse primer used for knockout of *bepA* | This study |
| cpxR-KO-F | ATTCAGGCTGCAAACATGCGTCAG | Forward primer used for knockout of *cpxR* | This study |
| cpxR-KO-R | GAATCGAGCTTGGGTAACATCAAAACCAAC | Reverse primer used for knockout of *cpxR* | This study |
| csgC-KO-F | AGGTTGGCTTTGGTAACAACGCG | Forward primer used for knockout of *csgC* | This study |
| csgC-KO-R | CTTGAGGGTTGTGTTATCCATACTTTCAGG | Reverse primer used for knockout of *csgC* | This study |
| csgE-KO-F | ACGGTTAAAACGCATCTTTATAATCTTTT | Forward primer used for knockout of *csgE* | This study |
| csgE-KO-R | TTACCACCAAAGTTTGGATTACGG | Reverse primer used for knockout of *csgE* | This study |
| csgG-KO-F | TGACAGATCGTAAAACCGGACAAACCTCG | Forward primer used for knockout of *csgG* | This study |
| csgG-KO-R | AATCAAAAATAAAGCGTGAGGGGCACTCAC | Reverse primer used for knockout of *csgG* | This study |
| dacA-KO-F | ACGTTAACAAAGTCACGCGGAAAGTCAG | Forward primer used for knockout of *dacA* | This study |
| dacA-KO-R | GGCGTAACGTAATTAAATAGCAACTCCCG | Reverse primer used for knockout of *dacA* | This study |
| degP-KO-F | TTTTACCTTTTGCAGAAACTTTAGTTCG | Forward primer used for knockout of *degP* | This study |
| degP-KO-R | TGCAAATGCCTAAAGGATGATGAAG | Reverse primer used for knockout of *degP* | This study |
| degQ-KO-F | CATTTAATCTGGTGTCTCATTGTTAGCCG | Forward primer used for knockout of *degQ* | This study |
| degQ-KO-R | TTCACAAACATGATGGAGGCGTC | Reverse primer used for knockout of *degQ* | This study |
| nlpC-KO-F | CGAATAATCCATTTACGGAATTTTTGTCTG | Forward primer used for knockout of *nlpC* | This study |
| nlpC-KO-R | GAATGCTGATTTCGACCATCTGATTTC | Reverse primer used for knockout of *nlpC* | This study |
| nlpI-KO-F | TTTTAACCGGGCAGGACGCCTTGTTAG | Forward primer used for knockout of *nlpI* | This study |

| | | | |
|---|---|---|---|
| nlpI-KO-R | AGGTGTATTAACGAACAACAACGCCCTCAC | Reverse primer used for knockout of *nlpI* | This study |
| prc-KO-F | CAGGCACAATTTCTTGTGCCTGATTGATATTAC TTGAGGGAGCGGGTTGGTGTAGGCTGGAGCTG CTTC | Forward primer used for knockout of *prc* | This study |
| prc-KO-R | TGTGCGCGCAGAACACCTGGTGTTCTGAAACG GAGGCCGGGCCAGGCATGCATATGAATATCCT CCTTA | Reverse primer used for knockout of *prc* | This study |
| ptr-KO-F | CATAGGCTGACTCGTGCAGCACAAG | Forward primer used for knockout of *ptr* | This study |
| ptr-KO-R | GCCGCAATCGTAAAGGTTTTGCCTG | Reverse primer used for knockout of *ptr* | This study |
| rcsB-KO-F | CATCTGATTCGTGAGAAGGATGTTCCAGG | Forward primer used for knockout of *rcsB* | This study |
| rcsB-KO-R | TAATGGGAATCGTAGGCCGGATAAGGC | Reverse primer used for knockout of *rcsB* | This study |
| spy-KO-F | CGATCCTGTCCACCGCATTACTG | Forward primer used for knockout of *spy* | This study |
| spy-KO-R | GTTAATCTCGTCAACACGGCACG | Reverse primer used for knockout of *spy* | This study |
| ydcP-KO-F | TTCCCTCATCCATTACCCGCGCAGTTAAG | Forward primer used for knockout of *ydcP* | This study |
| ydcP-KO-R | CAACGTTGAGTCATTTGATGCCGGATG | Reverse primer used for knockout of *ydcP* | This study |
| ydgD-KO-F | TCTGATTTTACTCCCACTTTCATCCCGTCC | Forward primer used for knockout of *ydgD* | This study |
| ydgD-KO-R | TAAAGGCTGGATTGGCCTGGTCTTGC | Reverse primer used for knockout of *ydgD* | This study |
| yhbU-KO-F | CTGCCTTAAATCAAAAATTGTCGCAGCAAG | Forward primer used for knockout of *yhbU* | This study |
| yhbU-KO-R | TGGCGGCCTGCTGATAAAATTCTTCC | Reverse primer used for knockout of *yhbU* | This study |
| DegP-His-pCA24N-F | GAGAAATTAACTATGAAAAAAACCACATTAGC ACTGAGTGCACTGG | Forward primer used for cloning of *degP-His* in pCA24N | This study |
| DegP-His-pCA24N-R | TGGCTGCAGGTCGAC*TTA*ATGATGATGATGATG A TGCTGCATTAACAGGTAGATGGTGCTGTC (Underline: $His_6$-tag; Italic: Stop codon) | Reverse primer used for cloning of *degP-His* in pCA24N | This study |
| DegP-S210A-F | ATCAACCGTGGTAACgctGGTGGTGCGCTGGTTA ACCTGAACGG (Small letters: Mutation sites) | Forward primer used for introduction of the point mutation (S210A) in *degP* | This study |
| DegP-S210A-R | AACCAGCGCACCACCagcGTTACCACGGTTGAT CGCTGCATCGGTC (Small letters: Mutation sites) | Reverse primer used for introduction of the point mutation (S210A) in *degP* | This study |
| DegQ-His-pCA24N-F | GAGAAATTAACTATGAAAAAACAAACCCAGCT GTTGAGTGC | Forward primer used for cloning of *degQ-His* in pCA24N | This study |
| DegQ-His-pCA24N-R | TGGCTGCAGGTCGAC*TTA*ATGATGATGATGATG ATGACGCATCAGCAGATAGATGCTTTCATTGC (Underline: $His_6$-tag; Italic: Stop codon) | Reverse primer used for cloning of *degQ-His* in pCA24N | This study |
| DegQ-S187A-F | ATTAACCGCGGTAACgctGGCGGTGCACTATTA AACCTTAACGGTGAG (Small letters: Mutation sites) | Forward primer used for introduction of the point mutation (S187A) in *degQ* | This study |
| DegQ-S187A-R | TAATAGTGCACCGCCagcGTTACCGCGGTTAAT GGAAGCATCTGTC (Small letters: Mutation sites) | Reverse primer used for introduction of the point mutation (S187A) in *degQ* | This study |
| Prc-His-pCA24N-F | GAGAAATTAACTATGAACATGTTTTTAGGCTT ACCGCGTTAGCTGG | Forward primer used for cloning of *prc-His* in pCA24N | This study |
| Prc-His-pCA24N-R | TGGCTGCAGGTCGAC*TTA*ATGATGATGATGATG ATGCTTGACGGGAGCGGGTTGTTCC (Underline: $His_6$-tag; Italic: Stop codon) | Reverse primer used for cloning of *prc-His* in pCA24N | This study |

| | | | |
|---|---|---|---|
| Prc-S452A-F | GACCGCTTCAGTGCTgctGCTTCAGAAATCTTTG<br>CCGCGGCAATG<br>(Small letters: Mutation sites) | Forward primer used for introduction of the point mutation (S452A) in *prc* | This study |
| Prc-S452A-R | AAAGATTTCTGAAGCagcAGCACTGAAGCGGTC<br>AACCAGCACCAC<br>(Small letters: Mutation sites) | Reverse primer used for introduction of the point mutation (S452A) in *prc* | This study |
| YdgD-pCA24N-F | GAGAAATTAACTATGCGTACAACCATTGCTGT<br>AG TGTTGGGTGC | Forward primer used for cloning of *ydgD* in pCA24N | This study |
| YdgD-pCA24N-R | TGGCTGCAGGTCGAC*TTA*TTTTTGCGACAGTTG<br>ATCCAGCTTGTCG<br>(Italic: Stop codon) | Reverse primer used for cloning of *ydgD* in pCA24N | This study |
| CsgA-sfGFP-pCold-F | GAAGGTAGGCATATGTCGAAAGGCGAAGAACT<br>GTTTACCGG | Forward primer used for cloning of *csgA-sfgfp* in pCold I | This study |
| CsgA-sfGFP-pCold-R | AGAGATTACCTATCT*TTA*TTTATACAGCTCATC<br>CATACCATGCGTAATCCC<br>(Italic: Stop codon) | Reverse primer used for cloning of *csgA-sfgfp* in pCold I | This study |
| CpxP-pCold-F | GAAGGTAGGCATATGGCTGAAGTCGGTTCAGG<br>CGATAACTG | Forward primer used for cloning of *cpxP* in pCold I | This study |
| CpxP-pCold-R | AGAGATTACCTATCT*CTA*CTGGGAACGTGAGTT<br>GCTACTACTCAATAGC<br>(Italic: Stop codon) | Reverse primer used for cloning of *cxpP* in pCold I | This study |
| YjfN-pCold-F | GAAGGTAGGCATATGCAGTCTGCAGAATTCGC<br>CAGTGCTGACTG | Forward primer used for cloning of *yjfN* in pCold I | This study |
| YjfN-pCold-R | AGAGATTACCTATCT*TCA*TGCATACAGTATCGC<br>CTGTACGCGCC<br>(Italic: Stop codon) | Reverse primer used for cloning of *yjfN* in pCold I | This study |
| mCherry-pCold-F | GAAGGTAGGCATATGAAGGGCGAGGAGGATA<br>ACATGGCC | Forward primer used for cloning of *mcherry* in pCold I | This study |
| mCherry-pCold-R | AGAGATTACCTATCTCTTGTACAGCTCGTCCAT<br>GCCGC | Reverse primer used for cloning of *mecherry* in pCold I | This study |
| sfGFP-pCold-F | GAAGGTAGGCATATGTCGAAAGGCGAAGAACT<br>GTTTACCGG | Forward primer used for cloning of *sfgfp* in pCold I | This study |
| sfGFP-pCold-R | AGAGATTACCTATCT*TTA*TTTATACAGCTCATC<br>CATACCATGCGTAATCCC<br>(Italic: Stop codon) | Reverse primer used for cloning of *sfgfp* in pCold I | This study |
| His-CsgA-pET28-Nde-F | CATATGGGTGTTGTTCCTCAGTACGG<br>(Underline: *Nde*I site) | Forward primer used for cloning of *csgA* in pET-28b | This study |
| His-CsgA-pET28-Bam-R | GGATCC*TTA*GTACTGATGAGCGGTC<br>(Italic: Stop codon; Underline: *Bam*HI site) | Reverse primer used for cloning of *csgA* in pET-28b | This study |
| Inverse-pET28-F | ACTGGTGGACAGCAAATGGGTCGGG | Forward primer used for linearization of pET-28b | This study |
| Inverse-pET28-R | CATGGTATATCTCCTTCTTAAAGTTAAACAAAA<br>TTATTTCTAGAGGGG | Reverse primer used for linearization of pET-28b | This study |
| CsgA-His-pET28-F | AGGAGATATACCATGGGTGTTGTTCCTCAGTAC<br>GGCGG | Forward primer used for cloning of *csgA* in pET-28b | This study |
| CsgA-His-pET28-R | TTGCTGTCCACCAGTGTACTGATGAGCGGTCGC<br>GTTGTTACC | Reverse primer used for cloning of *csgA* in pET-28b | This study |
| CsgA-A130G/S131G-F | GACCAGACTggcggcAACTCCTCCGTCAACGTGA<br>CTCAG<br>(Small letters: Mutation sites) | Forward primer used for introduction of the point mutations (A130G/S131G) in *csgA* | This study |
| CsgA-A130G/S131G-R | GGAGGAGTTgccgccAGTCTGGTCAACTGCAGCA<br>CC<br>(Small letters: Mutation sites) | Reverse primer used for introduction of the point mutations (A130G/S131G) in *csgA* | This study |
| RT-csgA-F | TCTGGCAGGTGTTGTTCCTC | Forward primer used for qRT-PCR of *csgA* | Sugimoto et al. 2018 (*13*) |
| RT-csgA-R | CCACCACCATGCTGGGTAAT | Reverse primer used for qRT-PCR of *csgA* | Sugimoto et al. 2018 (*13*) |
| RT-csgD-F | TCTCGTTATTAGACGCGCCG | Forward primer used for qRT-PCR of *csgD* | Sugimoto et al. 2018 (*13*) |

| | | | |
|---|---|---|---|
| RT-csgD-R | CAATGGATTGCAAGGCGTCC | Reverse primer used for qRT-PCR of *csgD* | Sugimoto et al. 2018 (*13*) |
| RT-ftsZ-F | CAATGGATTGCAAGGCGTCC | Forward primer used for qRT-PCR of *ftsZ* | Sugimoto et al. 2018 (*13*) |
| RT-ftsZ-R | TCAACACCTTCAATGCGCTC | Reverse primer used for qRT-PCR of *ftsZ* | Sugimoto et al. 2018 (*13*) |
